# CarboTag: a modular approach for live and functional imaging of plant cell walls

**DOI:** 10.1101/2024.07.05.597952

**Authors:** Maarten Besten, Milan Hendriksz, Lucile Michels, Bénédicte Charrier, Elwira Smakowska-Luzan, Dolf Weijers, Jan Willem Borst, Joris Sprakel

**Affiliations:** Laboratory of Biochemistry, Wageningen University and Research, Stippeneng 4, 6708 WE, Wageningen, the Netherlands; Institute of Functional Genomics in Lyon (IGFL), UMR5242, ENS de Lyon, CNRS, UCBL, 32-34 Avenue Tony Garnier, 68007 Lyon, France

## Abstract

Plant cells are contained inside a rigid network of cell walls. Cell walls are highly dynamic structures that act both as a structural material and as a hub for a wide range of signaling processes. Despite its crucial role in all aspects of the plant life cycle, live dynamical imaging of the cell wall and its functional properties has remained challenging. Here, we introduce CarboTag, a modular toolbox for live functional imaging of plant walls. CarboTag relies on a small molecular motif, a pyridine boronic acid, that targets its cargo to the cell wall, is non-toxic and ensures rapid tissue permeation. We designed a suite of cell wall imaging probes based on CarboTag in any desired color for multiplexing. Moreover, we created new functional reporters for live quantitative imaging of key cell wall features: network porosity, cell wall pH and the presence of reactive oxygen species. CarboTag opens the way to dynamical and quantitative mapping of cell wall responses at subcellular resolution.

## Introduction

Plant cells are characterized by the presence of a carbohydrate-rich cell wall. The cell wall plays a key role in a vast diversity of processes in the life cycle and survival of plants. It provides mechanical support to plant cells and tissues, regulates and modulates cell-to-cell communication and functions as a signaling platform for an enormous diversity of developmental and defense pathways [1–5]. From a physical perspective, plant cell walls are highly complex materials, consisting of structured interpenetrating networks of various carbohydrates, interspersed with a range of glycoproteins that can adaptively tune the mechanics and function of the wall, and contain numerous receptors for both chemical and physical signals [6–8]. Moreover, given its critical role in plant life, the maintenance of cell wall integrity, and its dynamic adaptation to meet environmental challenges or growth demands, are characterized by high genetic redundancy and many compensatory mechanisms [4, 9]. As a result, studying the cell wall has proven to be a formidable challenge, especially to monitor these highly dynamic processes in real-time in living plants and non-invasively. Yet, this information is crucial to study precisely what causes cell wall responses and their amplitude and localization, and what are its consequences.

As in all areas of cell biology, fluorescence microscopy is an invaluable tool in the study of plant cell walls. A wide variety of fluorescent probes have been reported to bind different cell wall components, recently reviewed comprehensively in [10]. These include the widely used fluorophores calcofluor white (cellulose) [11, 12], propidium iodide (PI, pectin) [13–15], aniline blue (callose) [16–18], basic fuchsin (lignin) [19, 20] and fluorol yellow (suberin) [21–23]. A range of antibodies for specific cell wall epitopes have been developed [24, 25]. However, these require fixation of the tissue and digestion of the cell wall to allow the bulky antibodies to permeate into tissues, and hence are not suitable for live imaging. Likewise, any of the chemical require fixation and tissue permeabilization for probe tissue permeation. Moreover, these probes and antibodies resolve localization, but do not provide functional insights about cell wall properties and their dynamics. To this end, a variety of genetically encoded cell wall biosensors for plants, in particular for the model plant *Arabidopsis thaliana*, are reported, for example, to reveal apoplastic calcium, glutamate encoded cell wall biosensors for plants, in particular for the model plant *Arabidopsis thaliana*, have been developed, for example to reveal apoplastic calcium, glutamate and pH levels [26–29]. Such genetically encoded probes require a genetic transformation of the species of interest, which is not possible for many organisms in the green lineage.

Despite significant advances in the chemical community to develop bespoke and functional probes for live imaging of animal cells [30, 31], efforts to develop de-novo probes tailored to plants, and to the cell wall in particular, are virtually absent. A key challenge in the de-novo design of probes for cell wall imaging is to target a fluorescent probe, or fluorescent reporter for some functional property of interest, specifically to the cell wall network. Cell wall targeting invariably involves binding of the probe to an epitope in the cell wall network. In addition, probes suitable for live cell imaging must be non-toxic and permeate specifically the apoplastic space of multicellular plant tissues without requiring a permeabilization or fixation step. Ideally, such an approach does not require genetic manipulation of the plant. One option is metabolic labeling, which has been established for root cell walls in *Arabidopsis thaliana* [32, 33], but this requires performing chemical reactions on the plant tissue, whose potential toxicity has not been investigated. Rather, we aimed to find a more convenient targeting strategy, encoded in the probe itself that could be applied to all plants without requiring growth on media supplemented with expensive chemically modified sugars. Previously, we used the pectin-binding peptide from extensin proteins to target a functional probe to the cell wall in live tissue [34, 35]. While successful, this approach is costly and resulted in low yields, the probes were unstable during prolonged storage, and showed relatively slow tissue penetration due to the relatively large hydrodynamic size of the peptide. We thus set out to find a reliable chemical motif for targeting molecules of interest to the cell wall.

Here we describe the design of a modular dye toolbox for the live and functional fluorescence imaging of plant cell walls. It is centered around a de-novo cell wall targeting motif, CarboTag, a small synthetic molecule that undergoes high-affinity binding with diols in the plant cell wall. CarboTag is non-toxic, even upon prolonged exposure, ensures rapid penetration into plant tissues and can deliver a range of water-soluble cargos to the cell wall. CarboTag allowed us to construct a range of cell wall fluorophores across the visible spectrum, based on readily available probes using click chemistry. Applying the same approach, we constructed chemical reporters for functional imaging of cell wall porosity, pH levels and Reactive Oxygen Species (ROS). CarboTag offers a modular approach for the specific delivery of a range of cargos to the plant cell wall and opens the way for creating functional and dynamical maps of living plant cell wall networks with subcellular resolution.

## Results

### CarboTag: a cell wall “zip code”

A generic molecular zip code that can address molecules fused to the cell wall has not yet been identified. We began our investigation by searching for a molecular motif that can target any fluorescent cargo to the cell wall reliably and with high affinity, which does not show cytotoxicity and ensures rapid permeation of the molecule into the apoplastic space of plant tissues.

We took inspiration from the chemistry used by plants themselves for cell wall binding. Plants use the natural compound boric acid B(OH)3, which is used as a crosslinker for cell wall components [36]. Boric acid can bind cell walls based on the dynamic covalent bonding that a boronic acid, RB(OH)2, undergoes with diols present in saccharides, including cell wall carbohydrates. For a typical boronic acid or phenylboronic acid, the optimal pH to form a diol complex lies between 7 and 9 [37]. As plant cell walls are generally acidic, with apoplastic pH levels ranging from 4 to 8 under physiological conditions [38], these motifs would only form very weak complexes with cell wall carbohydrates. By contrast, pyridinium boronic acids have been reported to form stable complexes in the correct acidic pH window [39, 40]. Starting from commercially available 4-pyridinylboronic acid and propargyl bromide, we synthesized, in a two-step reaction, the structure we name CarboTag (Fig.1a): a pyridinium boronic acid carrying a clickable alkyne group, suitable for conjugation to azide-functional fluorophores, or other azide-carrying cargo, using click chemistry [41–43].

Next, we synthesized a fusion of CarboTag to the photostable and water-soluble fluorophore AlexaFluor488 to give the CarboTag-AF488 probe. Live *A. thaliana* seedlings were stained in a solution containing 40 μM of CarboTag-AF488 for 30 mins and showed excellent staining of the root cell wall network, even deep inside the tissue (Fig. 1b). To verify that the CarboTag motif is responsible for the cell wall localization, as a control we incubated seedlings with the unmodified AF488 probe at the same concentration, which did not result in any detectable cell wall staining (Supplementary Fig.3). We also confirmed that CarboTag delivers the probe to the cell wall only, and does not result in membrane insertion, by performing a plasmolysis experiment on a CarboTag-AF488 stained seedling. Plasmolyzed roots show highly stained cell walls, and a complete lack of fluorescence signal from the plasma membrane (Supplementary Fig.9c,d).

**Figure 1.**
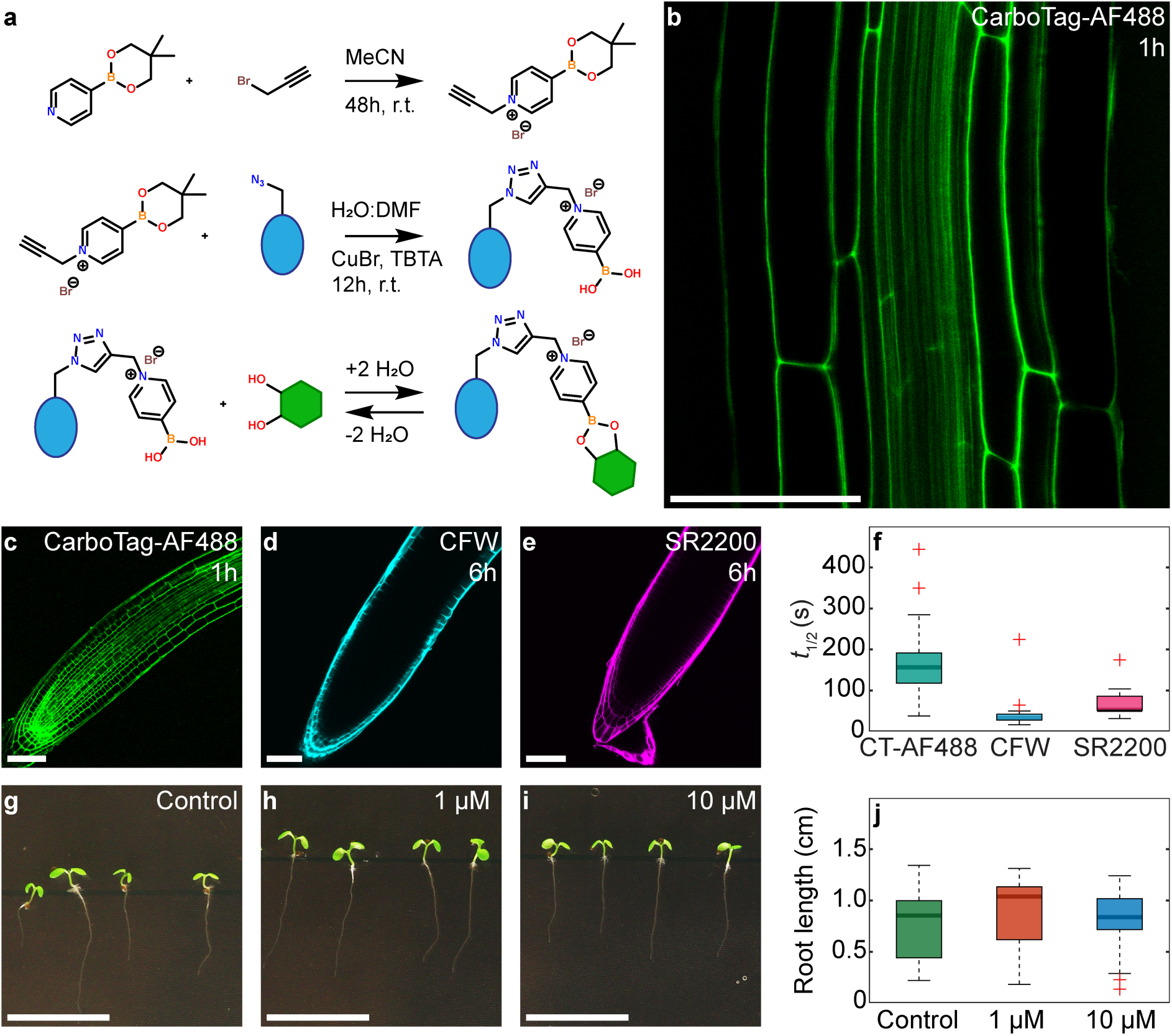
CarboTag cell wall targeting. a) Chemical route to synthesize CarboTag (top), CarboTag-modified dyes (middle) and chemical mechanism of forming reversible covalent bonds with diols (bottom). b) A. thaliana root stained with CarboTag-AF488. c-e) Arabidopsis roots stained with CarboTag-AF488, Calcofluor White (CFW) and Renaissance 2200 (SR2200), respectively, with incubation times indicated in the panels. f) Timescale of fluorescence recovery after photobleaching (FRAP), performed in Arabidopsis root hairs, for CarboTag-AF488 (CT-AF488), Calcofluor White and Renaissance 2200. g-j) Toxicity test of CarboTag, where Arabidopsis seeds are germinated and grown on agar media containing two CarboTag concentrations (as indicated), shown 5 days after germination (g-i) and corresponding measured root lengths (j). Scale bars represent 50 µm (b-e) or 1 cm (g-i)

Next, we benchmarked the performance of the CarboTag cell wall stain to the current state-of-the-art. We compared tissue permeation kinetics of CarboTag-AF488 with that of calcofluor white (CFW), Renaissance SR2200 and propidium iodide (PI), three commonly used cell wall stains [10], in live Arabidopsis seedling roots. Both CFW and SR2200 brightly stain the root epidermis after several hours but their penetration in deeper tissue is limited (Fig. 1d,e). We observe a similar lack of staining with PI when using isotonic 0.5 MS medium; PI staining improved significantly when using pure water as the staining medium, which is however hypotonic and thus imposed an osmotic stress on the tissue. Moreover, prolonged staining with PI, within 1h, leads to internalization of the dye and staining of the nucleus, and we observe severe signs of toxicity resulting from PI staining (Supplementary Fig.8).

By contrast, CarboTag-AF488 fully penetrates the root in as little as 15-30 mins and reaching full staining intensity in 1h (Fig.1b,c & S7). After 24h, CFW and SR2200 only had a marginal increase in penetration (Supplementary Fig.7). More importantly, roots exposed to CFW and SR2200 for 24h began to display signs of cytotoxicity, such as aberrant cell swelling (Supplementary Fig.7q,r), which are absent for CarboTag-AF488 (Supplementary Fig.7p). The de-novo CarboTag-based cell wall probe thus permeates tissues more rapidly and efficiently compared to the state-of-the-art cell wall probes.

The more rapid permeation of CarboTag compared to CFW and SR2200 could hint at a lower binding affinity to its cell wall epitope. To test if a weaker binding is responsible for the more rapid tissue permeation of our dye, we performed Fluorescence Recovery After Photobleaching (FRAP) experiments on these three probes in *A. thaliana* root hairs that were stained well by all probes. From the FRAP measurements, we extracted the characteristic timescale for fluorescence recovery and used this as a relative measure of the diffusion rate of the probes in the apoplastic space [44–46]. Both CFW and SR2200 diffuse more rapidly, indicated by a low recovery timescale, when compared to CarboTag-AF488, which has a slower fluorescence recovery (Fig.1f). The diffusion rate of probes in the apoplast is determined by their hydrodynamic size and their (un)binding kinetics to their wall epitope [47]. As the probes have similar sizes, the difference in diffusion kinetics suggests that CarboTag binds the cell wall more strongly compared to CFW and SR2200. This is expected given the strong dynamic covalent bonding between the pyridinium boronic acid and diols [37]. Thus, probe diffusion rates cannot explain the marked improvement in tissue permeation CarboTag offers. Both CFW and SR2200 are net negatively charged; the cell wall also has a net negative charge due to galacturonic acid residues [48]. Possibly, electrostatic repulsion plays a role in suppressing cell wall permeation. CarboTag-AF488 is zwitterionic, and thus net neutral, and shows relatively rapid permeation into the apoplastic network, despite binding the cell wall strongly.

To establish that CarboTag is non-toxic, we germinated and grew *Arabidopsis thaliana* seeds on agar plates containing 1 and 10 μM CarboTag. We compared their growth to seedlings germinated and grown on plates lacking the CarboTag addition to the medium. Seedlings exposed to both concentrations of CarboTag for 5 days show no visible aberrations and have identical root growth kinetics as compared to the control (Fig.1g-j).

### Multicolor plant cell wall imaging

Existing cell wall fluorophores encode their cell wall binding and fluorescence properties within the same chemical structure. As a result, there is only limited spectral flexibility in choosing a cell wall probe, which can be challenging in multiplexing experiments with plants that express a genetically encoded fluorescent reporter. The options are limited, especially for live mount imaging deep inside plant tissues, where many probes, such as CFW and SR2200, fail to penetrate well and PI leads to toxicity and cell internalization.

CarboTag can turn any azide-carrying and water-soluble fluorophore into a cell wall stain. Advances in dye chemistry have resulted in an enormous diversity of fluorophores that can be commercially procured with an azide modification. To demonstrate that CarboTag is compatible with a range of fluorophore chemistries, we selected probes across the visible spectrum based on coumarin (AF430), sulfo-rhodamine (AF488) and cyanine (sulfo-Cy3 and sulfo-Cy5) chemistries. Despite their very different chemical nature (see Supplementary Fig.1), all CarboTag fusions of these probes stain the cell walls of *A. thaliana* seedlings up to the vascular tissue within 1 hour (Fig. 2a-c). Access to a wide spectral range of cell wall stains facilitates multiplexed live imaging with existing plant lines that express genetically encoded fluorescent markers, such as for ABD2-mCherry (Actin; Fig. 2j) or MAP65-RFP (Microtubules; Fig. 2k).

**Figure 2.**
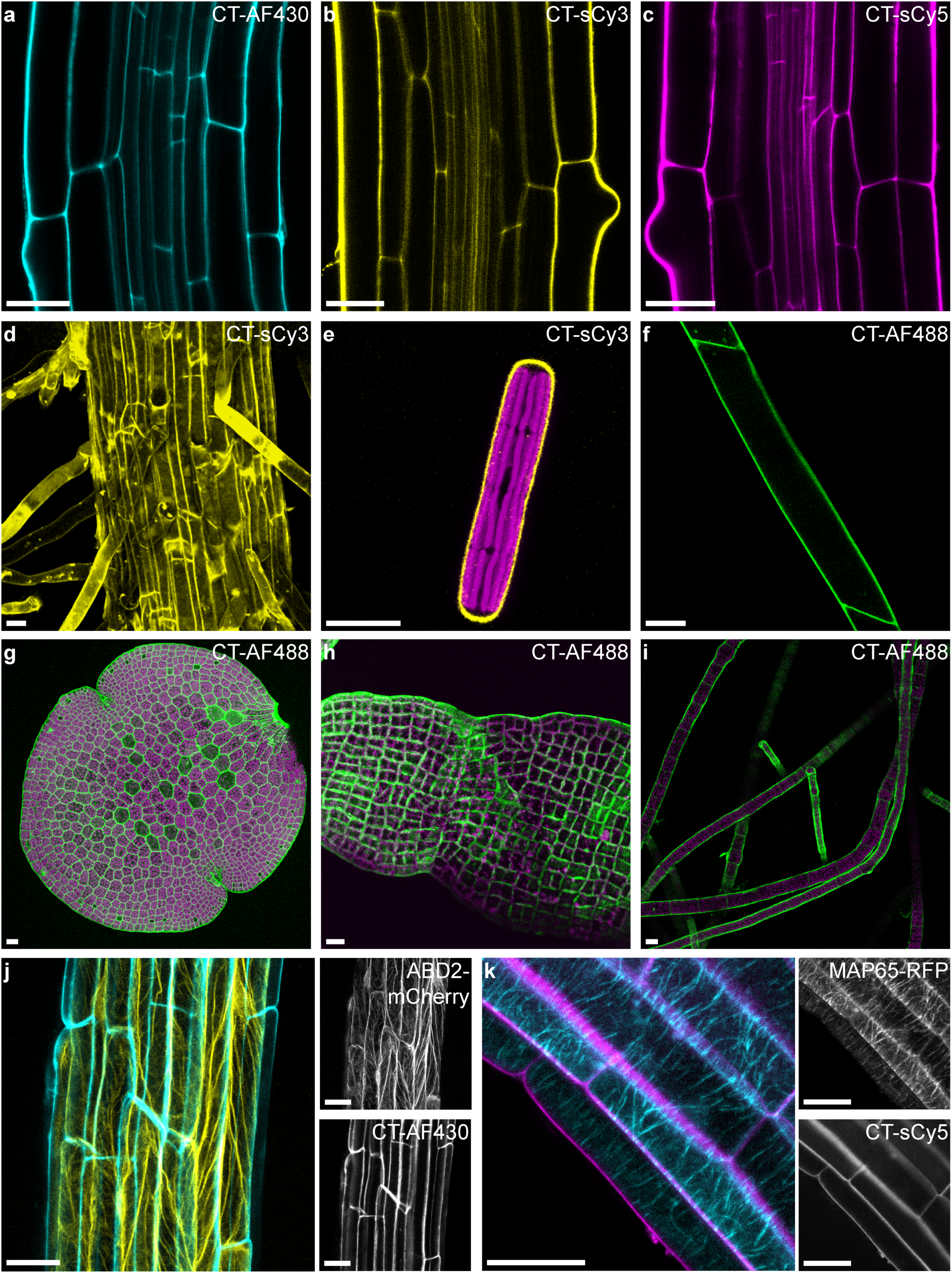
Multicolor CarboTag dyes function in plants and algae the green lineage: a-c) *A. thaliana* roots stained with CarboTag-modified AF430 (a), sulfo-Cy3 (b) and sulfo-Cy5 (c) that cover the emission spectrum from blue/green to far red. d) Root of the fern *C. richardii* stained with CarboTag-sCy3. e) single algal cell (*P. margaritaceum*) stained with CarboTag-sCy3, chloroplasts autofluorescence in magenta. f) Caulonema of the moss *P. patens* stained with CarboTag-AF488. g) Gemma of the liverwort *M. polymorpha* stained with CarboTag-AF488, chloroplast autofluorescence in magenta. CarboTag-AF488 staining of brown algae: *S. latissima* (h) and *S. rigidula* (i), chloroplast autofluorescence in magenta. CarboTag dyes for multiplexed imaging; j) CarboTag-AF430 (cyan) with actin marker ABD2-mCherry (yellow) and k) CarboTag-sCy5 (magenta) with microtubule marker MAP65-RFP (cyan), both in Arabidopsis root tissues. Scale bars in all images represent 25 µm.

To explore the compatibility of CarboTag with different plants along the green lineage, we performed staining experiments in a range of plant species: a green alga (*Penium margaritaceum*), a fern (*Ceratopteris richardii*), a moss (*Physcomitrium patens*) and a liverwort (*Marchantia polymorpha*). The cell walls of all these species could be stained with CarboTag-based probes (Fig.2d-g), indicating a broad applicability of CarboTag in different plant species. Moreover, it indicates that the epitope that CarboTag binds is present in all plant species and in green algae, even though their cell wall composition varies substantially [49–53].

We also tested the staining of organisms outside the plant lineage that feature a saccharide-rich cell wall. Interestingly, we found that CarboTag could *not* stain the cell wall of bacteria, both gram positive (*Bacillus subtilis*) and negative (*Escherichia coli*), nor oomycetes (*Phytophthora palmivora*). Clearly, CarboTag does not generally bind all diols, as there are many diol-containing polysaccharides and glycans present in the cell walls of these species. Since the oomycete cell wall contains cellulose [54, 55], this compound is not the CarboTag target. Thus, pectin is likely the epitope that is bound in green plants, as pectin is missing in the organisms we could not stain. Interestingly, we then tested if CarboTag could target cell walls of two species of brown algae; both *Saccharina latissimi* (Fig. 2h) and *Sphacelaria rigidula* (Fig. 2i) showed excellent staining with CarboTag-AF488, with the cell outlines clearly visible. The cell wall of brown algae lacks pectin and is mostly composed of alginates. Moreover, we observed that CarboTag stained agar-containing media and gels. We thus conclude that CarboTag binds hydrated carbohydrate hydrogels consisting of flexible polymers (pectin, agar, alginate), where the diol groups are readily accessible for binding.

### Cell wall porosity: CarboTag-BDP

The primary cell wall of plant cells is a complex composite material that must sustain enormous internal turgor pressures while also accommodating cell expansion and growth or tissue remodeling [2, 6, 56, 57]. This requires cell walls to be mechanically dynamic and plastic structures. We previously reported that a phenyl-BODIPY (BDP) molecular rotor probe, coupled to a pectin-binding peptide, could be used to probe changes and defects in cell wall network structure, which can affect their mechanical resilience [34, 35]. The BDP rotor, undergoes an intermolecular rotation upon photoexcitation to a non-radiative, dark, state. This physical motion requires fluid to flow around the molecule, coupling the molecular rotation hydrodynamically to its surroundings. This results in a BDP fluorescence lifetime that is low when the BODIPY probe rotates freely, e.g. in aqueous media, and high when the probe rotation is severely hindered [58, 59]. This feature allows the mapping of the mechanical environment of the dye with Fluorescence Lifetime Imaging (FLIM). In the Supplementary Information (Supplementary Fig.5) we show that the quantity that is probed is in fact the mesh size, or porosity, of the pectin network in the cell wall, and thus an indirect measure for the pectin levels in the cell wall space.

We coupled BDP to our CarboTag motif (see Fig 3a, b); the resulting probe CarboTag-BDP showed clear wall localization and rapid tissue permeation (Fig. 3c,f). To demonstrate its responsiveness to wall porosity in-planta, we manipulate the composition of the cell walls of Arabidopsis seedling roots by growing seedlings on a solid medium containing 3 nM of the cellulose synthase inhibitor isoxaben (ISX). Previous reports have shown that Arabidopsis seedlings grown on a solid medium containing 2.5 to 4.5 nM for 8 days showed limited growth and increased osmosensitivity [9, 60–62], while the levels of de-esterified pectin, and in some cases lignin, are found to be increased as a response mechanism to reduced cellulose synthesis. This leads to a denser meshwork in the cell wall upon cellulose synthase inhibition. [9, 63] We indeed find that cell walls in plants treated with isoxaben show a significantly higher lifetime of the CarboTag-BDP probe compared to both the control and mock-treated seedlings (Fig.3c -e). A higher lifetime indicates a more hindered intramolecular rotation, hence an increased density of the pectin network in the cell walls whose cellulose biosynthesis was inhibited.

**Figure 3.**
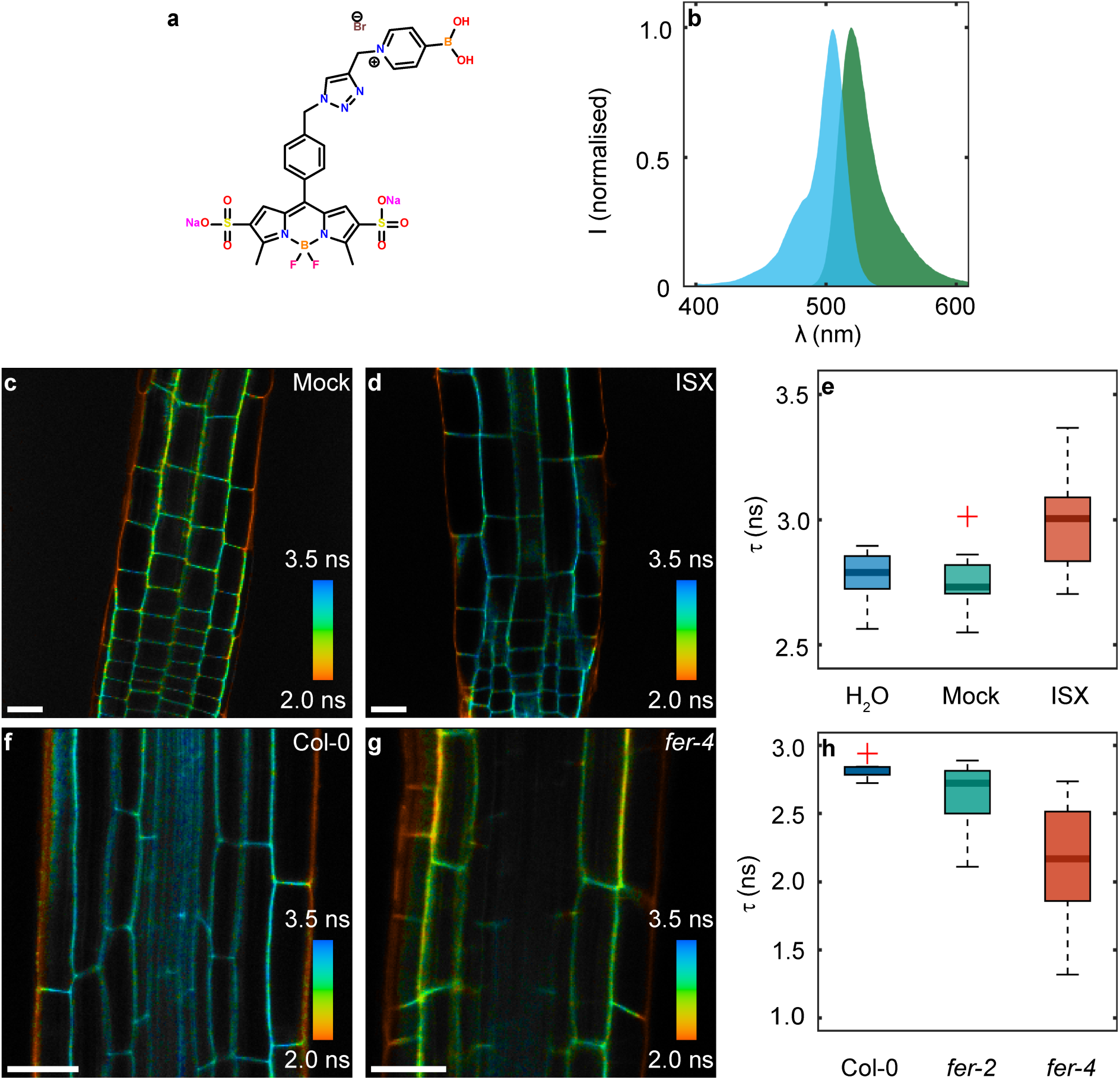
CarboTag-based cell wall porosity imaging. a) Chemical structure of CarboTag-BDP porosity probe. b) Normalized excitation (cyan) and emission (green) spectra of CarboTag-BDP c-d) Fluorescence lifetime imaging of Arabidopsis roots stained with CarboTag-BDP in wild type (Col-0, c) and FERONIA loss-of-function mutant fer-4 (d). e) Comparison of mean fluorescence lifetimes in the root cell walls, excluding the outer epidermal cell walls, in these lines including fer-2: n = 8 (Col0), n = 12 (fer-2) and n = 12 (fer-4). f-h) FLIM images of mock-treated (DMSO, f) and isoxaben (ISX, g) treated Arabidopsis seedling roots. h) Corresponding comparison of mean fluorescence lifetimes of cell walls in the root elongation zone excluding the epidermal cell wall: n = 8 (H2O), n = 11 (mock) and n = 18 (ISX). Color scale in FLIM images corresponds to the fluorescence lifetime. Scale bars in all images represent 25 µm.

As the cell wall is a vital structure to many processes, its integrity is continuously monitored by a complex cell wall integrity (CWI) maintenance pathway [4, 64]. A central player in CWI is FERONIA (*fer*), a pleiotropic pectin-binding receptor-like kinase [65–68]. Loss-of-function *fer* mutants show compromised CWI, resulting in accelerated root growth, spontaneous cell bursting and hypersensitivity to osmotic stress [66, 68, 69]. Interestingly, the phenotype can be rescued by embedding the tissue in a stiffer agar gel [66]. This suggests that FER mutants have weakened cell walls that have difficulties in sustaining internal turgor, unless supported by an exterior mechanical scaffold. However, AFM indentation experiments on FER mutants did not show a difference in the cell wall elasticity [68]. Since cell wall stiffness is dominated by cellulose microfibrils, FER mutants appear unaffected in cellulose levels and the origins of their mechanical problems lie elsewhere.

We performed FLIM-based porosity imaging of cell walls in *fer-2* and *fer-4* loss-of-function mutants stained with CarboTag-BDP. Mutant lines reveal significantly increased cell wall porosity, i.e., lower lifetimes than the wt control (Fig.3f-h). The *fer-4* line shows a more severely affected cell wall porosity, consistent with reports of a stronger root growth phenotype in *fer-4* versus *fer-2* [69]. Moreover, we find that the porosity of the *fer* mutants is significantly more heterogeneous compared to *wt*, indicated by the width of the lifetime distributions (Fig.3h). Given that our probe is sensitive to porosity at the nanometric scale, these changes likely reflect reduced pectin density and increased cell wall inhomogeneity in *fer* mutants.

### Apoplastic pH: CarboTag-OG

Plant cell growth requires that the rigid cell wall yield from the inflationary pressure of cellular turgor. In the “acid growth” theory, this is mediated by an acidification of the apoplast. A lower cell wall pH activates expansins, a family of non-enzymatic cell wall loosening proteins, resulting in cell walls with increased extensibility [70, 71],[72, 73]. The plant signaling molecule auxin controls growth by activating processes that acidify the apoplast, including the upregulation of proton pump synthesis and the activation of proton pumps by rapid phosphorylation through a non-canonical auxin signaling pathway [74–76]. Pathogens are also known to manipulate cell wall pH levels, most likely to loosen the cell wall and reduce the mechanical resistance to tissue colonization [77–79].

A commonly used approach to quantify apoplastic pH relies on the pH-sensitive ratiometric dye HPTS [80, 81]. However, HPTS is not only a pH reporter, but also a potent photo-acid, releasing protons upon illumination with the light required for imaging, thereby altering the pH as one tries to measure it [82]. HPTS illumination is even reported to result in strong cytotoxicity [83]. Alternatively, proton-sensitive dyes, such as the FS dye, are suspended in the medium to measure the pH at the surface of the plant tissue [74, 84]. These approaches measure pH at the surface of the root, thus detecting protons exuded from the plant. To measure the pH inside the cell wall, genetically encoded pH reporters for the cell wall have been described [29, 85]. All existing approaches have an intensity-based or ratiometric read-out, which makes the measurements sensitive to local changes in probe concentration and chromatic artefacts due to light scattering.

We use CarboTag to construct a FLIM-based apoplastic pH sensor, by coupling it to an Oregon Green (OG) fluorophore (Fig. 4a,b), with a reported pKa of 4.8. *In vitro*, the fluorescence lifetime of CarboTag-OG shows a distinct sigmoidal change with varying pH (Fig. 4c). The fact that OG can be used as a pH sensor with a lifetime-based read-out was reported previously [86] but not yet exploited in biological imaging. CarboTag-OG selectively targets the cell wall with the same staining kinetics (Fig. 4d-e).

**Figure 4.**
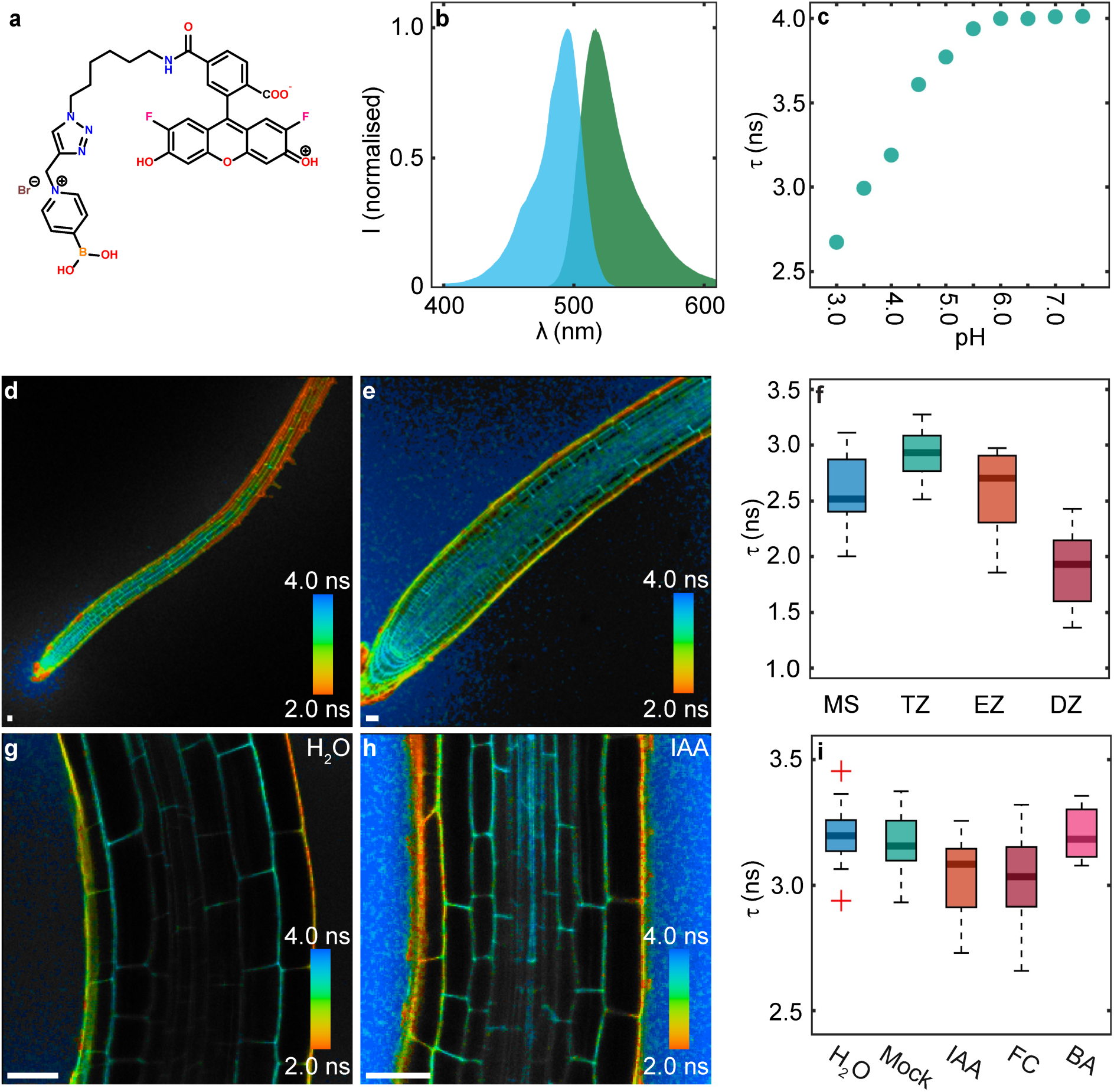
CarboTag based cell wall pH imaging. a) Chemical structure of CarboTag-OG pH probe. b) Normalized excitation (cyan) emission (green) spectrum of CarboTag-OG. c) Mean fluorescence lifetime of CarboTag-OG in 0.5x MS with different pH values. d-e), FLIM images of roots, f) Comparison of mean fluorescence lifetimes observed in the root meristem (MS, n = 10), transition zone (TZ, n = 10), elongation zone (EZ, n = 10) and differentiation zone (DZ, n = 10). g-h), FLIM images of non-treated roots (g) and roots treated with indole acetic acid (IAA) (h) after 10 min. j) Comparison of mean fluorescence lifetimes observed in the epidermis-cortex elongation zone cell wall between non-treated (n = 20), mock (DMSO, n = 12) treated, IAA (n = 17), fusicoccin (FC, n = 24) and benzoic acid (BA, n = 8) treated seedlings. Scale bars represent 25 µm.

FLIM map of roots of Arabidopsis seedlings shows that the fluorescence lifetime, and hence apoplastic pH, is lower in the elongation zone of the root compared to the root tip and maturation zone (Fig. 4d-f). This is consistent with acid growth theory where the apoplast is acidified in areas of the tissue where cells undergo expansion, and in agreement with previous reports of root surface pH patterns [74, 84]). We feel that it is important to note that the lifetime measurements performed in aqueous solutions of varying pH (Fig. 4c) cannot be used as a quantitative calibration curve, as the local chemical microenvironment in the apoplast, which influences the fluorescence lifetime of these probes, is far more complex than that in a simple buffer solution. This is, in general, true for comparing “calibration” curves constructed in simple solutions vs. using reporter probes in complex biological media. The pH-lifetime response curve can serve to evaluate the relative amplitude of pH variations within an experiment. We estimate, based on the data in Fig.4c, that the change of pH in the elongation zone, with respect to the root tip meristem is ΔpH = −1.1, while the change in pH between transition and elongation zone is ΔpH = −0.4. Clearly, the growth of the plant cells in the elongation zone is accompanied by a significant acidification.

Next, we subjected roots, pre-stained with CarboTag-OG, to treatment with 1 μM of the natural auxin Indole 3-Acetic Acid (IAA), which activates proton pumps through phosphorylation [74, 76] or 10 μM of fusicoccin, a fungal toxin that activates H+ ATPases [78]. Both treatments should lead to the efflux of protons from cells into the apoplast, resulting in rapid acidification. Indeed, the median fluorescence lifetime, analyzed in the epidermis-cortex cell wall, after 10 minutes of exposure to IAA or fusicoccin, is significantly lower compared to untreated control roots (Fig 4g-i). Based on the curve in Fig. 4i, we calculate the amplitude of the induced acidification to be ΛλpH = −0.22 for auxin treatment and ΛλpH = −0.31 for fusiccocin treatment. As a negative control, we subjected roots to treatment with benzoic acid (BA), a molecule with similar acidity and size to IAA but which lacks auxin’s biological activity, did not result in a decrease of the fluorescence lifetime.

### ROS generation in the cell wall: CarboTag-Ox

ROS, such as hydrogen peroxides and superoxide, are key signaling components in a large diversity of physiological processes ranging from development to wounding responses [87–90]. Reactive oxygen also plays an antimicrobial role in plant defenses, where ROS bursts are among the earliest responses of plant cells to pathogen recognition [90, 91]. Moreover, ROS acts as a chemical reagent, for example in crosslinking extensin glycoproteins in the cell wall [7, 92]. Various approaches, based on either colorimetric staining, oxidation-sensitive fluorophores or genetically-encoded biosensors, are reported to resolve ROS generation inside the cytosol, mitochondria or membranes of plant cells [93–97]. However, ROS in the apoplast itself has remained impossible to detect directly, while this is where ROS serves its function in crosslinking cell wall compounds and where it is delivered after pathogen perception. Moreover, the role of the apoplast in ROS perception is emerging through the putative oxidative modification of cysteine residues in the extracellular domains of CRRKs [98–102].

Our attempt to create an apoplastic ROS reporter for plants began by selecting the ROS reactive probe BODIPY 581/591 C11 (Fig. 5a). This probe features a BODIPY core, whose conjugated system is extended by a phenyl butadiene tail with two reactive sites for oxidation. In its native state, this extended pi-conjugated system emits in the red range. Upon oxidation of the tail, the conjugated system is shortened, shifting the emission to the green range. Due to its hydrophobic nature, this probe localizes to cellular plasma membranes and has been used for the ratiometric imaging of ROS production in mammalian and algal cells [103, 104].

**Figure 5.**
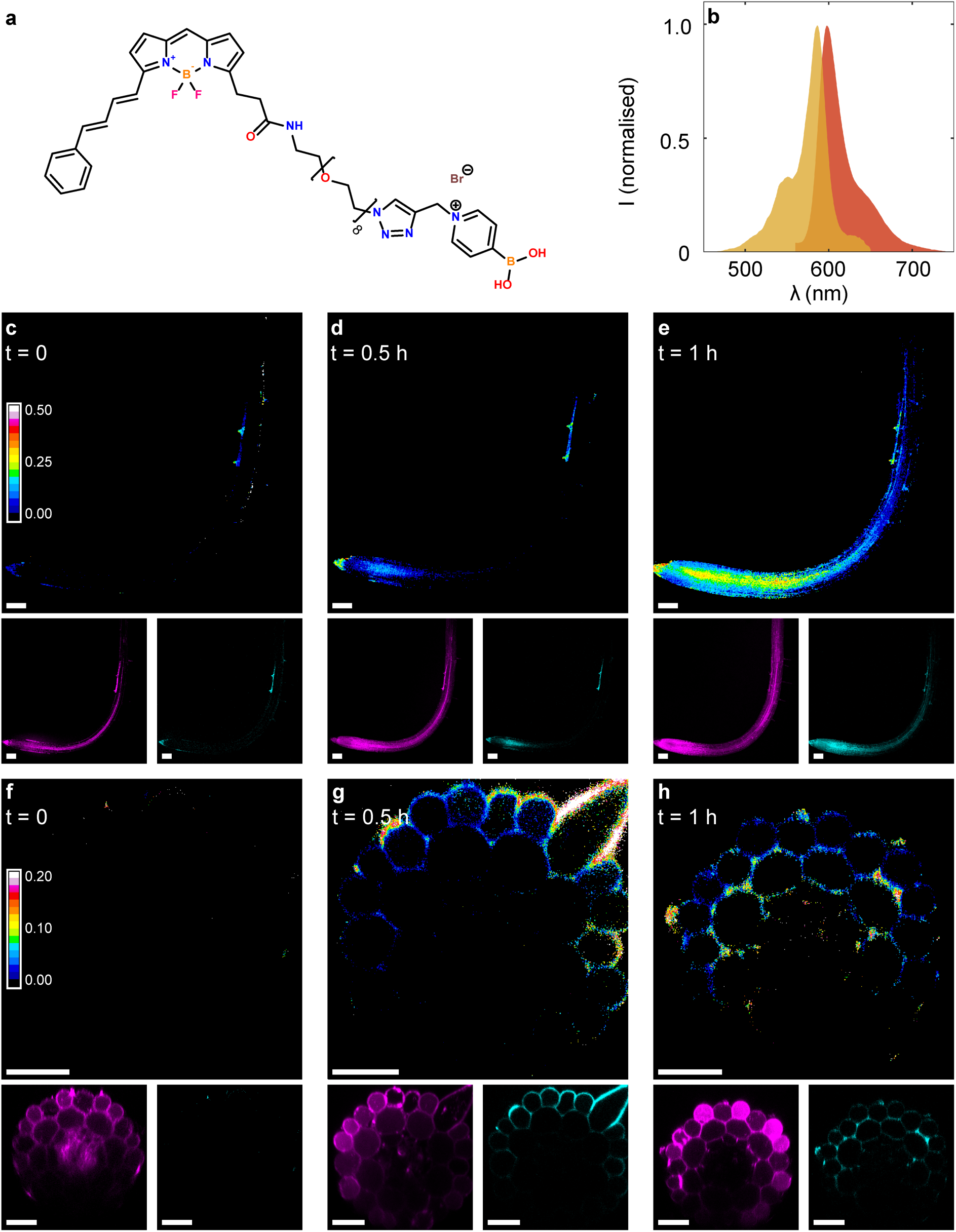
CarboTag-based cell wall ROS imaging. a) Chemical structure of CarboTag-Ox apoplastic ROS probe. Normalized excitation (yellow) and emission (orange) spectrum of non-oxidized CarboTag-Ox. c-e) Timelapse of CarboTag-Ox stained Arabidopsis roots treated with exogenous ROS, the color corresponds to the fraction of oxidized probe of the total fluorescence signal (oxidized + non-oxidized). Non-oxidized and oxidized signal are shown separately in magenta and cyan, respectively. f-g) Timelapse of CarboTag-Ox ROS probe stained Arabidopsis roots treated with flg22, color corresponds to the fraction of oxidized probe of the total fluorescence signal (oxidized + non-oxidized). Non-oxidized and oxidized signal are shown separately in magenta and cyan respectively. Scale bars represent 100 µm (c-e) and 25 µm (f-h) respectively.

Fusion of BODIPY C11 directly to CarboTag following the chemistry used above resulted in a probe that targets cell walls, but due to its amphiphilic nature also showed significant membrane insertion (Supplementary Fig.6a). To reduce membrane insertion, and thereby increase the specificity of cell wall localization, we introduced a hydrophilic spacer, consisting of a short oligoethylene oxide linker, between the BODIPY C11 probe and the CarboTag motif. This resulted in the probe we name CarboTag-Ox (Fig. 5a,b). This modification significantly reduced membrane insertion, although it did not completely prevent it (Supplementary Fig.6c). Based on the fluorescence intensity between membrane and cell wall following plasmolysis, we estimate that ∼70% of the signal in cells originates from the cell wall (Supplementary Fig.6e).

To validate functioning of CarboTag-Ox, we exposed pre-stained plants to radical oxygen species generated in the medium through a so-called Fenton’s reaction of iron ions, derived from hemin, with cumene hydroperoxide. Before ROS exposure, the probe emits purely in the red range, indicative of little to no oxidation, resulting in a red-to-green intensity ratio of close to zero (Fig. 5c). Over the course of an hour, we observed ROS activating the probe, first in the root tip (Fig. 5d) and spreading over the entire root (Fig. 5e), visible as an increasing red-to-green ratio, indicative of progressive oxidation of CarboTag-Ox. This shows that CarboTag-Ox can resolve the kinetics of ROS action in plant cell walls with live whole-mount imaging.

Next, we treated plants with the flagellin peptide *flg22*, which is perceived by the Arabidopsis FLS2 (Flagellin Sensitive 2) receptor as a signature of a potential pathogenic threat [105]. The resulting pattern-triggered immune (PTI) response involves the generation of a ROS burst, as a rapid and early defensive countermeasure [106]. Also here, before treatment, we observe little to no oxidation of the CarboTag-Ox probe (Fig. 5f). Treatment with *flg22* results in ROS production in the epidermis (Fig. 5g), which spreads as time proceeds to the cortical cells inside the root (Fig. 5h).

## Discussion

We presented CarboTag, a small and non-toxic cell wall targeting molecular motif as the basis for a modular toolbox for live and functional imaging of plant cell wall networks. In this work we showed how CarboTag can specifically localize any water-soluble fluorophore to the cell wall of various plants, which appears independent of the precise chemical make-up of the cargo.

### Advantages

As compared to the state-of-the-art, our CarboTag approach offers several advantages. The probes permeate tissues rapidly, without requiring potentially stressing permeabilization agents. They are non-toxic, while currently used stains for live-cell imaging of cell walls show severe toxicity phenotypes. Given the flexibility of the modular platform, cell wall stains can be constructed in any color or with any desired spectral features, which makes them ideal for multiplexing experiments. Moreover, they allowed us to construct probes for functional imaging of cell wall properties, that are applicable to a wide range of plants and brown algae species, without the need for genetic manipulation of the organism of interest.

### Limitations

The required incubation time for good tissue staining appears to be independent of the “cargo” that is attached to the targeting motif but is sensitive to the tissue type, plant species and dye concentration. At 40 μM, Arabidopsis seedling roots show staining of the vasculature in as little as 15 min, while at 10 μM, it may require up to an hour. Longer incubation can lead to increased dye uptake and signal intensity. While *A. thaliana roots,* fern roots and brown and green algae accept the stains rapidly (30-60 mins), some tissues and species were more challenging. Gemmae of *M. polymorpha* and thalli of *C. richardii* took up to 6h, or overnight, to stain at 40 μM. Lateral root primordia, before emergence, in Arabidopsis could not be stained at all with CarboTag-based probes. While the exact origins of reduced staining efficiency in these samples are unknown, we speculate that the presence of barrier layers, such as a thick cuticle on the surface of gemmae and thalli, could be responsible for the reduced uptake of these, by design, very hydrophilic dyes. The use of CarboTag-based probes in plant species and tissue types that we have not yet tested, especially those featuring cuticles or apoplastic barriers, may thus require optimization and could, in some cases, be challenging without the adding permeabilizing agents.

The FLIM-based reporters for cell wall porosity and pH are designed to be responsive to their local environment. As a result, these types of probes are generally solvatochromic, i.e., sensitive to their local chemical environment. This has several consequences for their use in *in-vivo* imaging. We observed that lifetime values for CarboTag-BDP and CarboTag-OG in the outer wall of root epidermal cells, at the root-medium surface, are consistently below the lowest lifetime for the probe in calibration media. This is indicative of probe quenching in these walls by chemical interaction with some, yet unidentified compound, which hinders analysis of the outer root surface. Similar effects were, in exploratory experiments, seen around tissue sites infected with pathogens or nematodes, where we believe they may be due to damage-associated compounds such as ROS or lignin precursors [107–109].

A second consequence of the solvatochromism of many lifetime-based fluorescent reporters is that their quantitative calibration *in vitro* is difficult. The chemical microenvironment in the complex plant cell wall is intrinsically very different from that in simple solutions, making a quantitative mapping from simple experiments to *in-vivo* results unreliable. For example, for the apoplastic pH probe CarboTag-OG, we performed lifetime measurements as a function of pH in aqueous solutions. Lifetime values of the same probes in-planta would suggest apoplastic pH levels that are significantly more acidic than those reported previously [80, 84]. Whether this is the result of solvatochromism due to the cell wall microenvironment that shifts the lifetime-pH curve, or due to inaccuracy in reported values, we cannot distinguish at this point. Out of caution, we have avoided converting our lifetime imaging results for this probe into absolute pH units and have used it as a tool to reveal relative acidification patterns and responses upon treatment. In general, we feel that great caution should be exercised in using calibrations for these types of probes performed in simple solutions to measurements performed in the complex environment of living cells.

### Outlook

We speculate that CarboTag can be used beyond what was reported here, and we envision extensions of this approach for functional mapping of other cell wall properties in the future, such CarboTag fusions to specific reporters for calcium, other ionic species or redox states. More hydrophobic cargo results in amphiphilic molecules that show a propensity for inserting into the membrane as well as binding the cell wall; this can be overcome to some extent by the introduction of a hydrophilic spacer, thereby extending the range of possibilities of this targeting strategy. One can also think beyond fluorescent probes and sensors, as CarboTag can theoretically also deliver bio-active signals into the cell wall to prevent their internalization in the cell; this could be a valuable tool to discriminate between the apoplastic and intercellular signal perception of, for example, hormones.

## Materials & Methods

### General

A complete overview of experimental details, including synthesis protocols, growth and preparation of biological material, imaging and analysis are provided in the Supplementary Information.

### General synthesis of CarboTag fluorophores

CarboTag fused fluorophores were prepared from commercially available azide functionalized dyes through copper-catalyzed azide alkyne cycloaddition (CuAAC). Coupling reactions take place in 1:2 water-dimethylformamide (DMF) mixtures with CuBr and tris(benzyltriazolylmethyl)amine (TBTA) as catalyst and ligand, respectively. After an overnight reaction, the resulting solution is diluted with water and filtered using a 0.45 μm syringe filter. The resulting solution is lyophilized and redissolved in water.

### Imaging

Imaging experiments were performed on a Leica TCS SP8 inverted confocal microscope coupled to a pulsed white light laser at a 40 MHz repetition rate, except for the experiments on brown algae which were performed on a Zeiss LSM880 confocal microscope. CarboTag-AF488, CarboTag-BDP (porosity probe) and CarboTag-OG were excited with a 488 nm line. Fluorescence was captured between 500 and 550 nm using a Hybrid detector. CarboTag-sCy3 and CarboTag-sCy5 were excited with a 548 nm and 646 nm lines, respectively. Fluorescence was detected between 558 and 608 nm (CarboTag-sCy3) and 656 and 706 nm (CarboTag-sCy5) using a Hybrid detector. CarboTag-Ox was simultaneously excited with a 488 nm and 581 laser lines, both at 5% laser power, to excite the oxidized and non-oxidized form of the probe respectively. Fluorescence was captured simultaneously between 500 and 550 nm for the oxidized form and between 591 and 641 nm for the non-oxidized form using Hybrid detectors in photon counting mode. MAP65-RFP was excited with a 561 nm laser line. Fluorescence was captured between 570 and 615 nm using a Hybrid detector. ABD2-mCherry was excited with a 585 nm laser line. Fluorescence was captured between 600 and 650 nm using a Hybrid detector. For CarboTag-AF430 we used a 440 nm diode pulsed laser (40 MHz) and fluorescence was captured between 500 and 550 nm. Fluorescence was captured through a 10x (NA 0.3) dry objective or a 63x (NA 1.2) water immersion objective depending on the sample type.

Ratiometric measurements performed with CarboTag-Ox were processed in ImageJ. For both the oxidized (green) and non-oxidized (red) channel background signal was removed. The pixel values of these 2 images were corrected for their backgrounds. Subsequently the resulting pixel values were added up and corrected pixel values of the green channel were divided by the pixel values of the combined image. The resulting 32-bit image has pixel values between 0 and 1 which indicate the fraction of total signal originating from the oxidized (green) channel. This fraction is displayed in a 16-color scale.

FLIM experiments using CarboTag-BDP and CarboTag-OG were performed on a Leica TCS SP8 inverted confocal microscope coupled to a Becker-Hickl SPC830 time-correlated single photon counting (TCSPC) module. An excitation line of 488 nm was used and fluorescence was captured between 500 and 550 nm using a hybrid detector. Acquisition times were between 60 s and 120 s depending on signal strength were used to obtain an image of 256 x 256 pixels. Resulting FLIM images were processed in SPCImage 8.5 software to obtain two-component exponential decay curves for every pixel. FLIM images are presented in a false-color scale that represents the mean fluorescence lifetime per pixel in nanoseconds.

### Comparison CT-AF488 vs CFW & SR2200

5-day old Arabidopsis seedlings were incubated in a 12 well plate in 0.5x MS with either 40 μM CarboTag-AF488, a 1000x diluted Renaissance 2200 (SR2200) stock solution (from supplier) or a 20x diluted CalcoFluor White (CFW) solution (from a 1g/L solution). The well plate was covered with and left on a plate shaker at 60 rpm. Seedlings were taken out after 15 minutes, 30 minutes, 60 minutes, 3 hours, 6 hours and 2 hours, washed with clen 0.5x MS and imaged. Imaging was performed on a Leica SP8 multiphoton system with a pulsed Coherent Chameleon Ti:sapphire laser. CarboTag-AF488 was excited with a 980 nm line, and the fluorescence was captured between 500 and 570 nm using a hybrid detector. Both SR2200 and CFW were excited with the 2-photon laser at 810 nmand the fluorescence was captured between 420 and 500 nm using a Hybrid detector. Fluorescence for all 3 dyes was captured through a 40x (NA = 1.10) water immersion objective.

### FRAP

5-day old Arabidopsis seedlings were incubated in 0.5x MS with either 40 μM CarboTag-AF488, a 1000x diluted Renaissance 2200 (SR2200) stock solution (from supplier) or a 20x diluted CalcoFluor White (CFW) solution (from a 1g/L solution). FRAP experiments were performed on a Nikon C2 CLSM with an argon laser using a 60x (NA =1.40) oil immersion objective. CarboTag-AF488 was excited with a 488 nm argon laser line, fluorescence was captured between 500 and 550 nm using a PMT detector. Both SR2200 and CFW were excited with a 405 nm line, fluorescence was detected between 417 and 477 nm using a PMT detector. FRAP was performed by selection of a bleach ROI and a reference ROI on root hairs with the same size for all measurements. A series of images were taken as a pre-bleach (3 images, 8 second interval) where the laser power and detector gain were balanced to have a minimal number of saturated pixels. Bleaching was performed by increasing the laser power to 80-100% depending on the sample for 5 sec interval in a small region-of-interest. Following bleaching, a time-lapse sequence was taken at a frame rate of 0.13 fps. The fluorescence recovery in the bleach spot was normalized and corrected against the reference region to correct for imaging-induced photobleaching. The resulting data was fitted with a single exponential fit in Matlab R2021b to extract the half-time of recovery by using the maximum intensity value after bleaching as the plateau value [110].

### Plant growth

*A. thaliana* wild-type, *fer*-2 and *fer*-4 mutants were surface sterilized by washing them in a 50% ethanol solution, followed by a wash in 70% ethanol and a 100% ethanol solution (5 minutes per solution). The sterilized seeds were dried for 1h before storage. Sterilized seeds were put on half strength (0.5x) MS plates and placed at 4 ⁰C overnight before placing them vertically under long day growth conditions (16 h light, 8 h dark) for 5 days. 5-day old seedlings were transferred to 0.5x liquid MS medium containing 40 μM of the CarboTag dyes, except for CarboTag-Ox which was kept a 10 μM. Seedlings were incubated for 1 h and washed in fresh 0.5x MS medium for 1 min and transferred to a glass slide with 0.5x MS medium before imaging. Osmotic shock treatments were performed by placing the seedlings on a glass slide with a 0.5 M mannitol solution instead of 0.5x MS medium

*A. C. richardii* spores (Hn-n strain) were sterilized and grown as previously describe in a Hettich MPC600 plant growth incubator set at 28°C, with 16 hours of 100 μmol m^–2^ s^–1^ [111, 112]. Plants were grown on 0.5x MS medium supplemented with 1% sucrose. Gametophytes were grown from spores and synchronized by imbibing the spores in the dark in water for >4 days. Sporophytes were obtained by flooding plates containing sexually mature gametophytes with water. Sporophytes used for imaging were three to four weeks old. Samples were incubated with 40 μM of CarboTag dye for 6 h in 0.5x MS and washed with 0.5x MS for 1 min before imaging.

*B. P. margaritaceum* NIES 217 cells were grown in Woods Hole Medium (WHM) for 15 days) at 20 °C with long-day conditions, 30 – 50 µmol/m2/s light and with continuous agitation (60 RPMs, 50ml flasks). Liquid cultures with an OD750 of 0.2-0.3 were used for imaging experiments. After incubating with 40 μM of CarboTag dye for 6 h cells were centrifuged at 500 rpm for 1 min. Cells were transferred to fresh 0.5x MS and imaged.

*C. S. latissima* fragmented gametophytes were inoculated in Provasoli enriched seawater (PES) under low light conditions (16 μmol m-2 s-1), a 12h:12h light-dark cycle at 13 ⁰C. Transfer material to normal light conditions (50 μmol m-2 s-1) after 6 days, PES medium was changed every 7 days. Samples were incubated with 10 μM CarboTag dye for 12h and 90min, washed 3x with sea water >30 min and imaged.

*D. S. rigidula* male gametophytes were cultivated in PES at 16 ⁰C with a 12h:12h light-dark cycle. Photon irradiation was 30 μmol m^-2^ s^-1^ (white light spectrum). Samples were incubated with 10 μM CarboTag dye for 12h and 90min, washed 3x with sea water >30 min and imaged.

*E. M. polymorpha* was grown on B5 medium under continuous light (40 μmol m^-2^ s^-1^) at 25 ⁰C. Gemma were isolated using a 200 μl pipet tip and incubated with 40 μM CarboTag dye in 0.5x MS for 6h and washed for 1 min in clean 0.5x MS before imaging.

*F. P. patens* was grown on BCDAT medium under continuous light (40 μmol m^-2^ s^-1^) at 25 ⁰C. Samples were incubated with 40 40 μM CarboTag dye, washed with 0.5x MS for 1 min and imaged.

### Toxicity test

The toxicity of the CarboTag was determined by supplementing melted 0.5x MS gel with water (control), 1 μM or 10 μM CarboTag from a 100 mM stock solution in water. Sterilized seeds were plated, their root length was measured 5 days after germination.

### Chemical treatments

*Isoxaben:* Isoxaben treated seedlings were generated by supplementing melted 0.5x MS gel with 3 nM isoxaben from a 5 mM stock in DMSO. Sterilized seeds were plated, and placed at 4 ⁰C overnight followed by putting the plates vertically under long day growth conditions for 5 days before staining and imaging. A mock treatment was performed by adding an equal amount of DMSO without isoxaben to melted 0.5x MS gel *Fusicoccin:* 5-day old seedlings were stained with 40 μM CarboTag-OG for 1h and transferred to a microscopy slide with unbuffered 0.5x MS supplemented with 10 μM fusicoccin from a 5 mM stock in DMSO. 5 x 2 min continuous FLIM measurements were performed immediately after. A mock treatment was performed by placing seedlings in unbuffered 0.5x MS with 0.2% DMSO during imaging.

*Auxin:* 5-day old seedlings were stained with 40 μM CarboTag-OG for 1h and transferred to a microscopy slide with unbuffered 0.5x MS supplemented with 1 μM auxin from a 5 mM stock in DMSO. 5 x 2 min continuous FLIM measurements were performed immediately after. A mock treatment was performed by placing seedlings in unbuffered 0.5x MS with 0.2% DMSO during imaging.

*Benzoic acid:* 5-day old seedlings were stained with 40 μM CarboTag-OG for 1h and transferred to a microscopy slide with unbuffered 0.5x MS supplemented with 1 μM benzoic acid from a 5 mM stock in DMSO. 5 x 2 min continuous FLIM measurements were performed immediately after.

*Flg22:* Seedlings were stained with 10 μM CarboTag-Ox for 1h, transferred to 5 ml 0.5x MS containing 1 μM flg22 and vacuum infiltrated for 5 min. Seedlings were removed from this mixture before imaging. Mock treatment was performed by placing seedlings in 0.5x MS under vacuum for 5 min.

*ROS:* Seedlings were stained with 10 μM CarboTag-Ox for 1h and transferred to a microscopy slide with freshly prepared 0.5x MS supplemented with 5 μM hemin from a 1 mM stock in DMSO and and 2 mM cumene peroxide from a 500 mM stock in ethanol. Images of single roots are taken with a 15-minute interval for 1 h.

## Acknowledgements

This work and J.S. and M.B. are funded by the European Research Council project Catch, project number 101000981. The work of B.C. is funded by the European Research Council - project ALTER e-GROW, project number 101055148. Views and opinions expresses are however those of the author(s) only and do not necessarily reflect those of the European Union or the European Research Council Executive Agency. Neither the European Union nor the granting authority can be held responsible for them. We gratefully acknowledge valuable discussions, suggestions, probe testing and the supply of plant material by Sergio Martin Ramirez, Sjoerd Woudenberg, Polet Carillo Carrasco, Martijn de Roij, Alice Malivert, Benoit Landrein, Julie Guerreiro, Peter Marhavy, Antoine Chevalier and Thorsten Hamann.

## Author contributions

M.B. and J.S. established the CarboTag concept; M.B., J.S., J.W.B. and D.W. designed the experiments and contributed to data interpretation; M.B. and M.H. designed and performed the synthesis of the CarboTag motif; M.B. synthesized all CarboTag based probes and performed all in-planta experiments; Imaging experiments on brown algae were performed by B.C.; L.M. performed the sulfoBDP lifetime-viscosity experiment; J.S., J.W.B. and D.W. supervised the project. M.B., J.S. and J.W.B. wrote the manuscript. All authors reviewed the paper.

