## Supplementary Information for "CarboTag: a modular approach for live and functional imaging of plant cell walls"

**Vendors chemicals, dyes and treatments**

All chemicals used for synthesis and treatments of biological material were purchased from Merck, TCI chemicals and AK Scientific. Fluorescent probes were purchased from Lumiprobe and Broadpharm. Silica (irregular, 40-63 $\mu$m particle size) and silica C18 (spherical, 20-45 $\mu$m, 100A) for flash and reverse phase chromatography respectively were purchased from Screening Devices, eluent mixture ratios are reported by volume. $^{1}$H, $^{13}$C and $^{19}$F NMR spectra were recorded on a Bruker Avance III 400 MHz spectrometer.

**Synthetic procedures CarboTag, fluorophore precursors and CarboTag modified fluorophores**

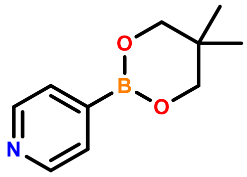


**Neopentyl glycol protected pyridine-4-boronic acid (1)**

Pyridine-4-boronic acid (2.4 g, 20 mmol) and neopentyl glycol (2.2 g, 20 mmol) were dissolved in dry toluene (250 ml). The mixture was heated to 130 $^{o}$C and refluxed overnight under $N_2$. The mixture was cooled to r.t. The formed precipitate was collected by vacuum filtration, washed with toluene and dried under vacuum to yield 2.44 g of neopentyl glycol protected pyridine-4-boronic acid. Yield: 64 %. $^{1}$**H NMR** (400 MHz, CDCl3) δ 8.73 – 8.42 (m, 2H), 7.79 – 7.44 (m, 2H), 3.74 (s, 4H), 1.02 (s, 6H). $^{13}$**C NMR** (101 MHz, CDCl3) δ 149.14, 128.20, 72.55, 32.05, 22.00.

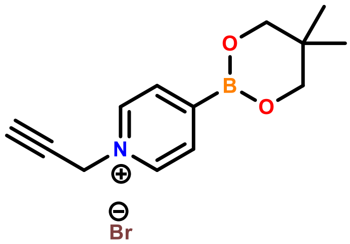


**CarboTag (2)**

Neopentyl glycol protected pyridine-4-boronic acid (764 mg, 4 mmol) was added to 100 ml acetonitrile. The mixture was purged with $N_2$ for 5 minutes before the dropwise addition of propargyl bromide (80% in toluene, 8.6 ml, 80 mmol). The mixture was left stirring for 48 h at r.t. under $N_2$. The precipitate was removed by filtration and filtrate was evaporated to dryness. The resulting brown solid was dried overnight to yield 989 mg of CarboTag. Yield: 80%.$^{1}$**H NMR** (400 MHz, D2O) δ 8.89 (d, J = 6.2 Hz, 2H), 8.26 (d, J = 6.2 Hz, 2H), 5.50 (d, J = 2.6 Hz, 2H), 3.39 (s, 5H), 3.28 (s, 1H), 0.86 (s, 7H). $^{13}$**C NMR** (101 MHz, D2O) δ 141.08, 130.60, 118.18, 79.13, 72.85, 67.27, 49.10, 35.15, 19.49.

**4(bromomethyl)-benzaldehyde (3)**

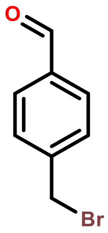


4(bromomethyl)benzonitrile (10 g , 51 mmol) was added to 100 ml of dry toluene and cooled to 0 °C in an ice bath. Add 72 ml DIBAL-H in hexane (1M, 72 mmol) over a period of 2h (6x12 ml) at 0 °C. The solution was left stirring for 1h at 0 °C before being diluted with 150 ml chloroform. 330 ml 10% HCl was slowly added, the mixture was allowed to heat up to r.t. and was left stirring overnight. The organic layer was isolated an washed with 3x 125 ml water, dried over $MgSO_4$ and concentrated to yield a white solid. The solid was recrystallized from hexane to yield 6.79 g of white, fine needle like crystalline solid. Yield: 67%. **$^1$H NMR** (400 MHz, CDCl3) δ 10.02 (s, 1H), 7.87 (d, J = 8.1 Hz, 2H), 7.56 (d, J = 7.7 Hz, 3H), 4.52 (s, 2H). **$^{13}$C NMR** (101 MHz, CDCl3) δ 191.64, 144.39, 136.29, 130.07 (d, J = 49.5 Hz), 32.09.

**4(azidomethyl)-benzaldehyde (4)**

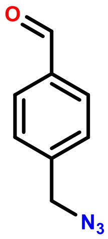


4-(bromomethyl)benzaldehyde (5g, 25 mmol) and $NaN_3$ (2.5 g, 38 mmol) were added to 50 mL of DMF in a round bottom flask. The mixture was stirred for 1.5 hours at 60 °C. After cooling the solution was diluted with 250 mL of ethyl acetate and washed with 2x250mL of water. The organic layer was dried with $MgSO_4$, filtered, concentrated and dried under vacuum to yield 4-(azidomethyl)benzaldehyde as an oil with pleasant smell (3.76 g, 93% yield). **$^1$H NMR** (400 MHz, CDCl3) δ 10.02 (s, 1H), 7.90 (d, J = 8.1 Hz, 2H), 7.49 (d, J = 8.5 Hz, 2H), 4.45 (s, 2H). **$^{13}$C NMR** (101 MHz, CDCl3) δ 191.74, 142.24, 136.30, 130.32, 128.59, 54.38.

**$N_3$-BODIPY porosity probe (5)**

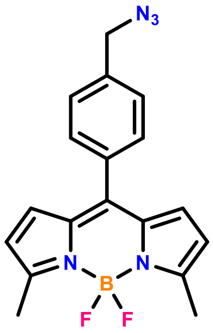


4-(azidomethyl)benzaldehyde (2.0 g, 12.41mmol) was added to a 1L 3-neck round bottom flask with 500 mL of anhydrous dichloromethane and 2-methylpyrrole (2.092 mL, 24.82 mmol). The solution was purged with $N_2$ for 30 minutes after which trifluoroacetic acid (500 $\mu$L, 6.2 mmol) was added dropwise. The mixture turned orange and was stirred for 2 hours after which 2,3-dichloro-5,6-dicyano-1,4-benzoquinone (2.82 g, 12.41 mmol) ws added followed by purging with $N_2$ for 10 minutes followed by 20 minutes of stirring. Di-isopropylethylamine (15.1 mL, 86.7 mmol) and boron trifluoride diethyl etherate (15.3 mL, 124 mmol) were added, the mixture was left stirring for 24 hours under $N_2$. The reaction mixture was diluted with 2:3 hexane:ethyl acetate followed by removal of dichloromethane under educed pressure. After purification on silica (2:3 hexane:ethyl acetate) the product was isolated as a red, crystalline solid (1.07 g, 25% yield). **$^1$H NMR** (400 MHz, CDCl3) δ 7.52 (d, J = 8.1 Hz, 2H), 7.43 (d, J = 8.1 Hz, 2H), 6.69 (d, J = 4.1 Hz, 2H), 6.27 (d, J = 4.1 Hz, 2H), 4.46 (s, 2H), 2.65 (s, 6H). **$^{19}$F NMR** (376 MHz, CDCl3) δ -147.61 (dd, J = 65.3, 32.1 Hz).

### Sulfonated $N_3$-BODIPY porosity probe (6)

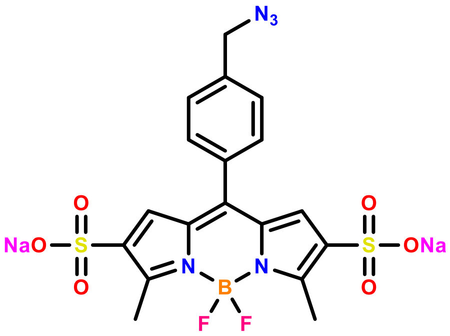


$N_3$-BODIPY (180 mg, 0.52 mmol) was dissolved in 24 ml of dry DMF after which sulfur trioxide pridine complex is added (1.632 g, 10 mmol). The mixture is purged with $N_2$ for 15 minutes and heated to 60 $^0$C for 24 hours after which it wascooled down to r.t. A saturated sodium carbonate solution was added until no more gas formation was observed. The resulting aqueous solution is washed with 2x 25 ml chloroform and concentrated. To remove the final traces of pyridine  40 ml of toluene is dded nd evaporated under eeduced pressure, this is repeated 3 times. The resulting solid is dissolved in absolute ethanol, filtered through a celite pad and the filtrate concentrated and dried under vacuum to yield 245 mg of sulfornated $N_3$-BODIPY with reasonable purity. Yield: 85%.**$^1$H NMR** (400 MHz, D2O) δ 7.59 (d, J = 5.3 Hz, 4H), 7.20 (s, 2H), 4.55 (s, 2H), 2.65 (d, J = 1.2 Hz, 6H).

**General synthesis protocol CarboTag modified fluorophores**

Prepare a 0.5 M CarboTag solution in MilliQ (MQ) water and a 50 mM tris[(1-benzyl-1H-1,2,3-triazol-4-yl)methyl]amine (TBTA) solution in DMF. Add copper(I) bromide (CuBr) (6 mg, 0.04 mmol) to a vial equipped with a stirring bar. Add 1 ml of TBTA solution (0.05 mmol) and stir for 5 min. Add additional DMF to ensure the final DMF:$H_2O$ ratio is 2:1 if needed. Add a concentrated solution of the fluorophore ($H_2O$ or DMF) and add 2 equivalents of CarboTag for every equivalent of dye. Add additional water to have a DMF:$H_2O$ ratio of 2:1. Seal with a septum and bubble with $N_2$ for 15 min, stir overnight at r.t. Dilute the solution with MQ water, filter out the precipitate with a 0.45 $\mu$m syringe filter. Freeze dry the filtered solution, redissolve in water and further dilute based on absorbance.

**Comment on general synthesis protocol**

Preparing separate solution of CarboTag, TBTA and the fluorophore allows for a simpler procedure since weighing milligrams these compounds dry is tedious and prone to error. Additionally, dissolving these compounds separately prevents precipitation and/or slow dissolving opposed to adding solvent to a mixture of dry reagents. Since copper(I) bromide and TBTA are both insoluble in water adding all DMF containing solutions before adding $H_2O$ prevents these compounds from precipitating. Their poor solubility is later exploited for purification by diluting the reaction mixture with water and removing the precipitate consisting of CuBr and TBTA while the desired product remains in solution. The quantity of CuBr and TBTA is kept constant independent of the amount of dye and reaction volume, at our largest scale synthesis (CarboTag-AF430) 25 mg of dye was used which amounts to a total of 0.032 mmol. This excess of catalyst and ligand makes changing their respective used quantities redundant.

**CarboTag-AF430**

The general synthesis protocol was followed, AF430-azide (25 mg, 0.032 mmol) was dissolved in 0.8 ml MQ water. 0.128 ml 0.5M (0.064 mmol) CarboTag solution was added. The total reaction volume was 3 ml.

**CarboTag-AF488**

The general synthesis protocol was followed, AF488-azide (20 mg, 0.029 mmol) was dissolved in 2 ml MQ water. 0.12 ml 0.5M (0.060 mmol) CarboTag solution was added. The total reaction volume was 6 ml.

**CarboTag-sCy3**

The general synthesis protocol was followed, sCy3-azide (10 mg, 0.0136 mmol) was dissolved in 1 ml MQ water. 0.054 ml 0.5M (0.0272 mmol) CarboTag solution was added. The total reaction volume was 3 ml.

**CarboTag-sCy5**

The general synthesis protocol was followed, sCy5-azide (10 mg, 0.0132 mmol) was dissolved in 1 ml MQ water. 0.052 ml 0.5M (0.0264 mmol) CarboTag solution was added. The total reaction volume was 3 ml.

**CarboTag-BDP**

Sulfonated N3-BODIPY porosity probe (50 mg, 0.09 mmol) and CarboTag (27.9 mg, 0.09 mmol) were dissolved in 2 ml water-DMF (1:3) and bubbled with $N_2$ for 5 min. TBTA (9.5 mg, 0.018 mmol), CsF (27.3 mg, 0.18 mmol) and CuBr (5.2 mg, 0.036 mmol) were added and the mixture was left stirring for 5 h. The mixture was concentrated under vacuum and purified on a C18 reverse phase silica flash column using a 5:95 to 20:80 methanol-water gradient.

**CarboTag-OG**

The general synthesis protocol was followed, OG-azide (5 mg, 0.009 mmol) was dissolved in 1 ml DMF. 0.036 ml 0.5M (0.018 mmol) CarboTag solution was added. The total reaction volume was 3 ml.

**CarboTag-Ox**

Azido-PEG$_8$-amine (8.6 mg, 0.0195 mmol), di-isopropylethylamine (2.5 mg, 0.195 mmol) and BDP581/591-NHS ester were added to 1 ml DMF, degassed with $N_2$ and left stirring overnight. 6 mg CuBr was added followed by 1 ml 50 mM TBTA in DMF. 0.52 ml 0.5 M CarboTag in MQ water was added followed by 0.95 ml MQ water. The reaction wasleft stirring overnight. After diluting with MQ water the resulting solution was filtered through a filter paper instead of a syringe filter.

**CarboTag-BDP581/591**

The general synthesis protocol was followed, BDP581/591-azide (5 mg, 0.01 mmol) was dissolved in 0.4 ml DMF. 0.04 ml 0.5M (0.02 mmol) CarboTag solution was added. The total reaction volume was 3 ml.

**<u>Spectroscopy</u>**

**Steady state fluorescence**

UV-Vis absorption measurements were recorded on a Shimadzu UV2600 Spectrophotometer using 1 cm path quartz cuvettes. Fluorescence emission and excitation measurements were performed on a Agilent Cary Eclipse Fluorescence spectrophotometer with a xenon lamp using 10 mm quartz cuvettes. Absorption of samples for fluorescence emission and excitation measurements were kept below 0.1.

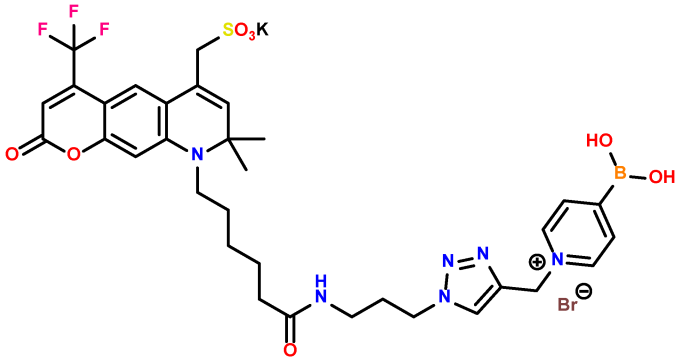

CarboTag-AF430

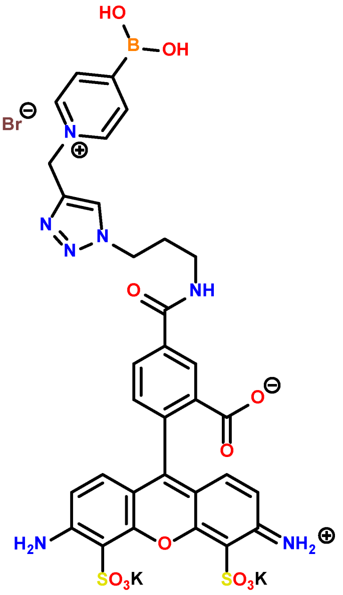

CarboTag-AF488

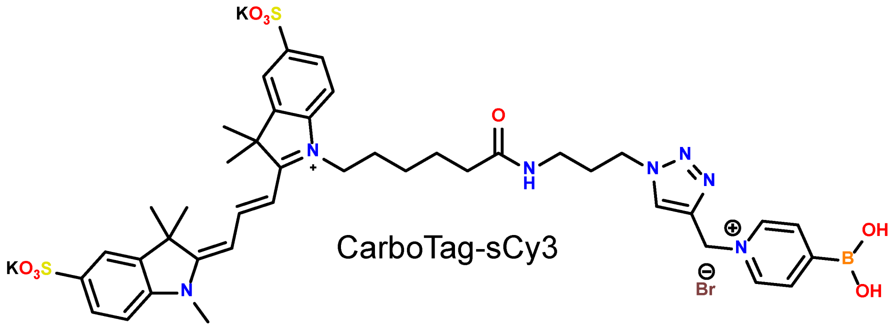

CarboTag-sCy3

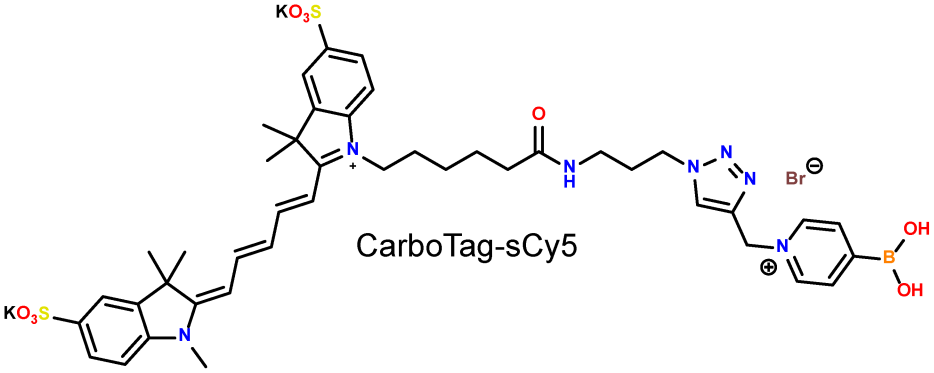

CarboTag-sCy5

**Figure S1:** chemical structures of commercially available azide functionalized fluorophores modified with CarboTag.

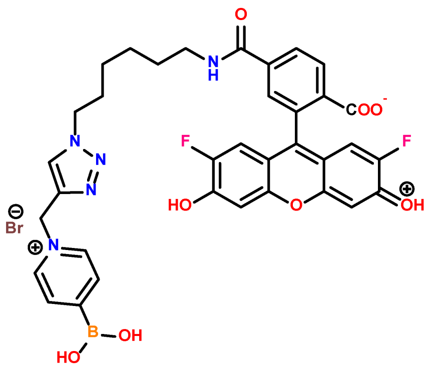

CarboTag-OG

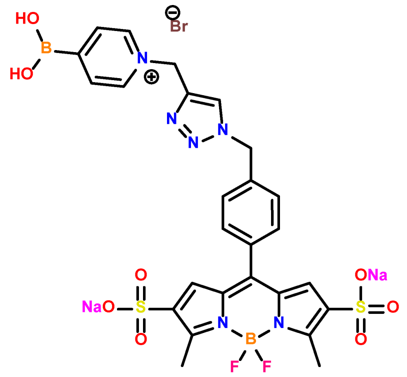

CarboTag-BDP

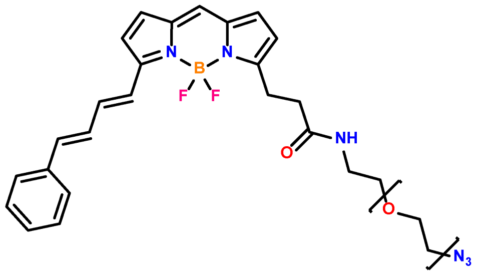

Pegylated ROS probe

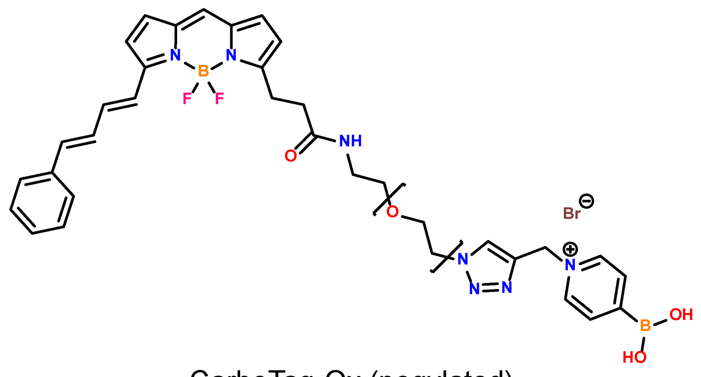

CarboTag-Ox (pegylated)

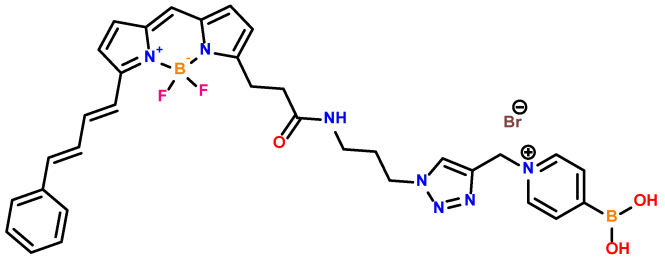

CarboTag ROS probe (non-pegylated)

**Figure S2:** chemical structures of pH (CarboTag-OG), porosity (CarboTag-BDP) and ROS (CarboTag-Ox) probes. A fraction of pegylated precursor of CarboTag-Ox (pegylated ROS probe) is likely to be present the isolated CarboTag-Ox. Similar pegylated probes have shown to internalize in the vacuole and cytoplasm over time which explains why there is some internalization of CarboTag-Ox in root cells. The membrane localizing CarboTag ROS probe lacking a PEG-8 moeity is shown in the bottom.

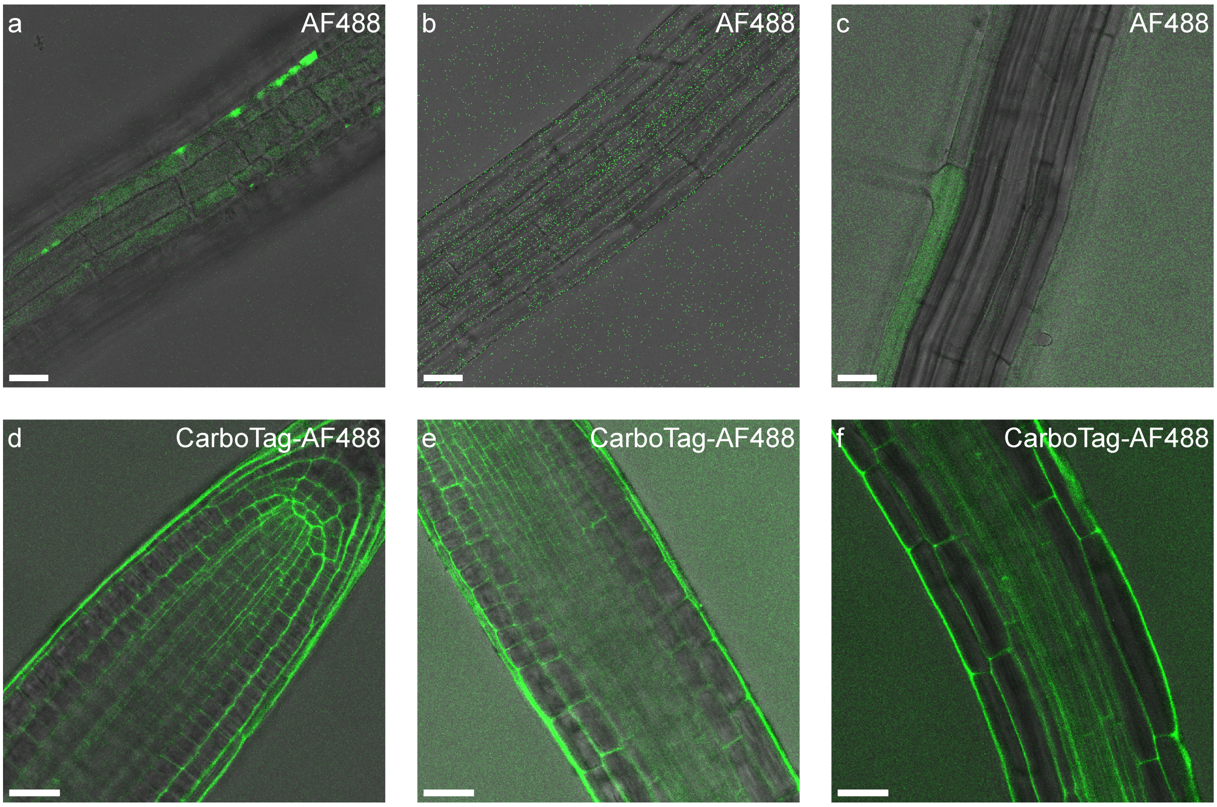


**Figure S3:** a-c) Arabidopis roots stained with non-functionalized Alexafluor488 (AF488) show limited staining. d-f) CarboTag-AF488 exhbitis superior staining over its non-modied counterpart. Scale bars represent 25 $\mu$m

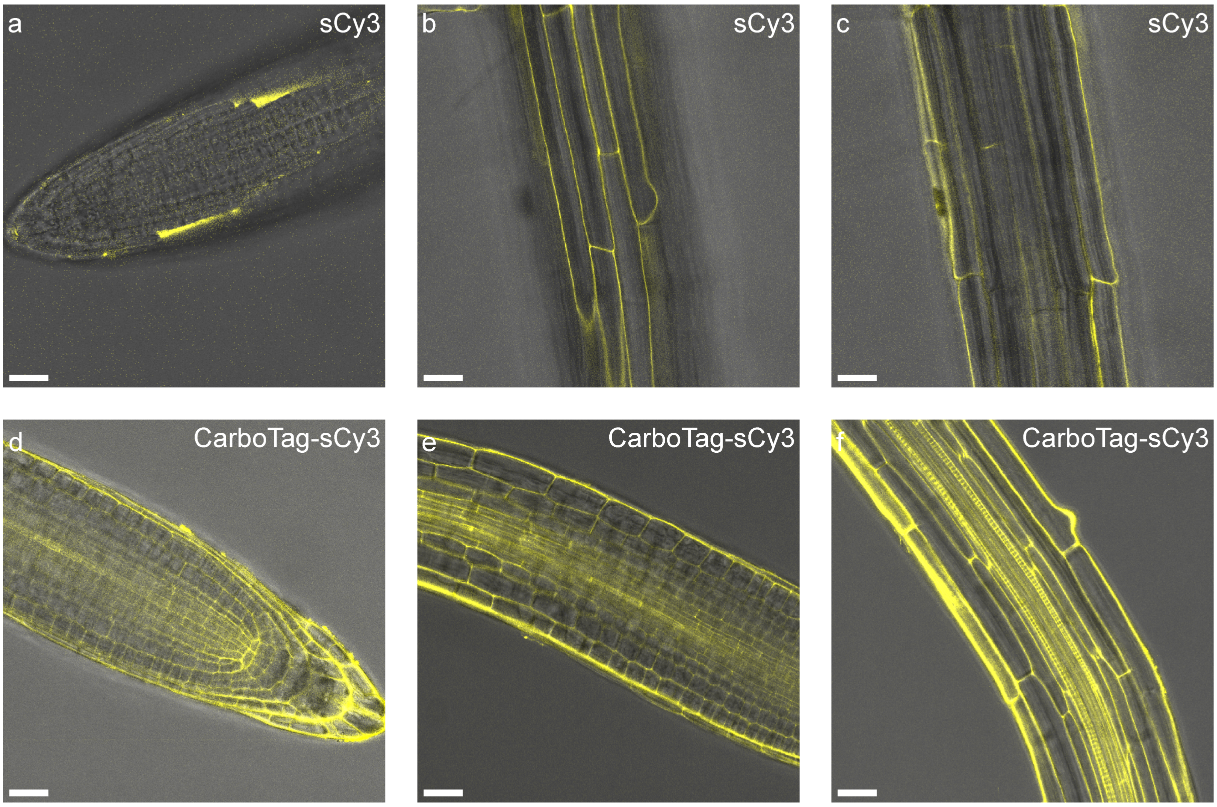


**Figure S4:** a-c) Arabidopis roots stained with non-functionalized sCy3 show limited staining. Non-specific staining of sCy3 in the root epidermis (b) is present but limited in deeper tissue (c). d-f) CarboTag-sCy3 and exhbitis superior staining over its non-modied counterpart. Scale bars represent 25 $\mu$m.

### MICRORHEOLOGY

We, and others, have used the term microviscosity to describe the physical quantity that these probes are sensitive to [35, 60, 61]. However, in the context of the plant cell wall this term is misleading. The probe binds a cell wall epitope and sits in the water-filled meshes in-between biopolymer chains; the local aqueous viscosity in these pores will not change substantially when the cell wall composition and structure are altered. Rather, we believe that cell wall targeted BDP probes map cell wall porosity [34, 35]. The flow field caused by probe rotation will extend out to several times the size of the molecule, i.e. approximately several to ten nanometers. In larger pores, e.g. cellulose rich maturated cell walls, the fluid can flow freely as it is not hindered by nearby polymer chains. In smaller pores, e.g. in pectin-rich young cell walls, the fluid flow is severely hindered by the network resulting in a slower rotation and hence a longer lifetime.

To test this hypothesis, we make use of the feature of polymer solutions that at the same mass concentration for a specific type of polymer, higher molecular weight polymers give a higher macroscopic solution viscosity but identical network mesh, or pore, size [62]. We measured the fluorescence lifetime of a sulfonated BDP rotor dissolved in solutions of the polymer poly(ethylene oxide) of different molecular weights and as a function of polymer concentration. For each sample, we independently measured the solution viscosity using light-scattering based microrheology [63, 64]. The BDP lifetime as a function of viscosity indeed does not yield a universal curve, indicating that the physical quantity that is measured is not the solution viscosity (Supplementary Fig. 5b). Rather, if we plot the fluorescence lifetime as a function of the mass concentration, we find a collapse of the data onto a single curve (Supplementary Fig. 5a); hence, the response is independent of polymer length and only of polymer concentration, i.e. mesh size of the polymer network. These results support the notion that BDP molecular rotors are cell wall porosity sensors.

We studied the fluorescence lifetime response of PEG-BDP in aqueous solutions of ethylene glycol, poly(ethylene glycol) of various chain lengths (Table S1) and poly(acrylic acid) 5.1kg $mol^{-1}$. The concentrations were varied between 0.01 g $cm^{-3}$ and 10 g $cm^{-3}$ until reaching the solute solubility limit. The viscosity of those solutions was determined by DLS micro rheology.

Table S1: The average molecular weight and polydispersity (PDI) of PEG polymers used to study the response of PEG-BDP to viscosity and mesh size.

| Polymer | $M_n$ (g $mol^{-1}$) | PDI |
|---|---|---|
| PEG1k | 1000 | <1.3 |
| PEG10k | 10000 | <1.3 |
| PEG100k | 95000 | 1.08 |
| PEG728k | 728000 | 1.24 |

PMMA-PEGMA particles with a hydrodynamic radius of 77 nm (PDI between 0.002 and 0.042) were synthesized by a standard emulsion polymerization.

In a 250 mL round-bottom flask, methyl methacrylate (8.9 g) and PEG methacrylate (0.5 g) were mixed in MQ water (100 mL) containing a small amount of SDS (20 mg). The mixture was degassed by purging with nitrogen for 10 min. The flask was put under slight vacuum. The biphasic mixture was stirred at 75°C for 15min, and the stirring speed adjusted so as to avoid emulsification. A solution of potassium persulfate (100 mg) in MQ water (5 mL) was injected, and the reaction carried out for at least 24h. The obtained turbid suspension was filtered through glass wool and stored at 4°C in a plastic bottle.

The dynamic light-scattering experiments were carried out on a Malvern Nano-S, with a He-Ne laser (λ=632.8 nm), an avalanche photodetector at a detection angle of 173°. In all experiments the temperature was controlled at 20°C. The DLS particles were suspended in the samples of interest by diluting the stock 1000 times. The scattered intensity autocorrelation function was measured 3 times, for 1000 s to 1 h depending on the characteristic decorrelation time of each individual sample. The mean square displacement of the particles was determined from the scattered intensity autocorrelation functions, and the diffusion coefficient computed by using the slope of the mean square displacement vs time plot at time scales beyond the caging dynamics. The diffusion coefficient was directly translated into a viscosity using the Stokes-Einstein equation.

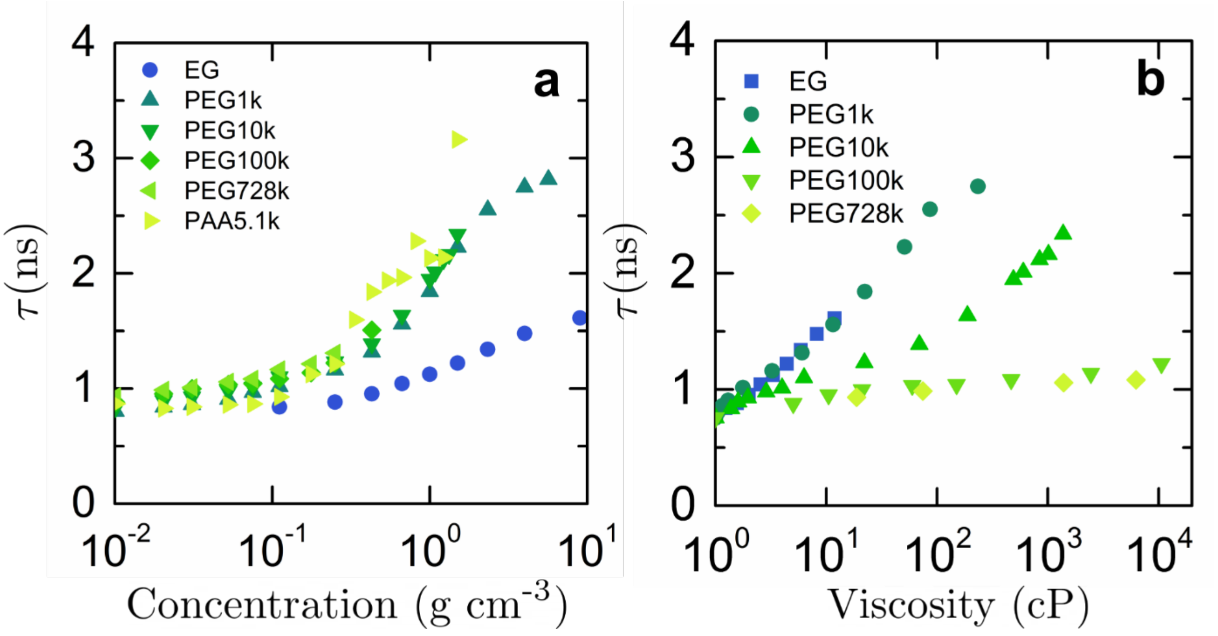


**Figure S5**: Evolution of the fluorescence lifetime of PEG-BDP in solutions of EG, PEG and PAA with various chain length, as a function of EG or polymer concentration (a), and as a function of viscosity as measured by DLS microrheology (b)

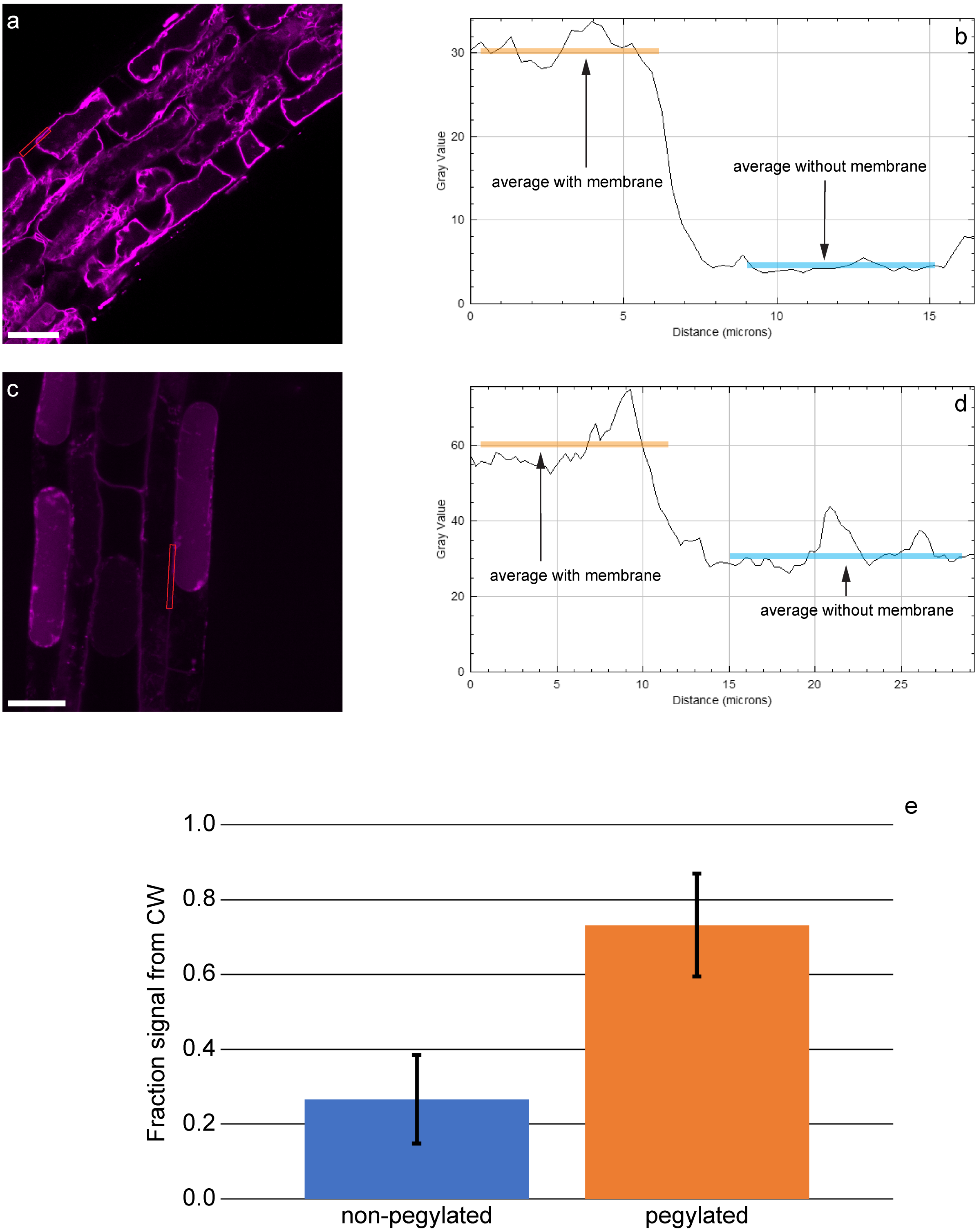


**Figure S6:** plasmolysis of non-pegylated CarboTag ROS probe (a) and pegylated CarboTag-Ox (c) shows high membrane affinity for the CarboTag ROS probe and limited membrane localization for CarboTag-Ox. CarboTag-Ox however internalizes which could potentially be caused by a pegylated intermdiate present in the staining solution. Using an ROI following the cell wall and membrane (in red) we determined the signal originating from the cell wall + membrane and only the cell wall for CarboTag ROS (b) and CarboTag-Ox (d). The fraction of the signal originating from the cell wall was calculated by dividing the average intensity in the cell wall by the average intensity of the area with both the cell wall and cell membrane (e), n=3 for the non-pegylated and n=4 for the pegylated probe. Scale bars represent 25 $\mu$m.

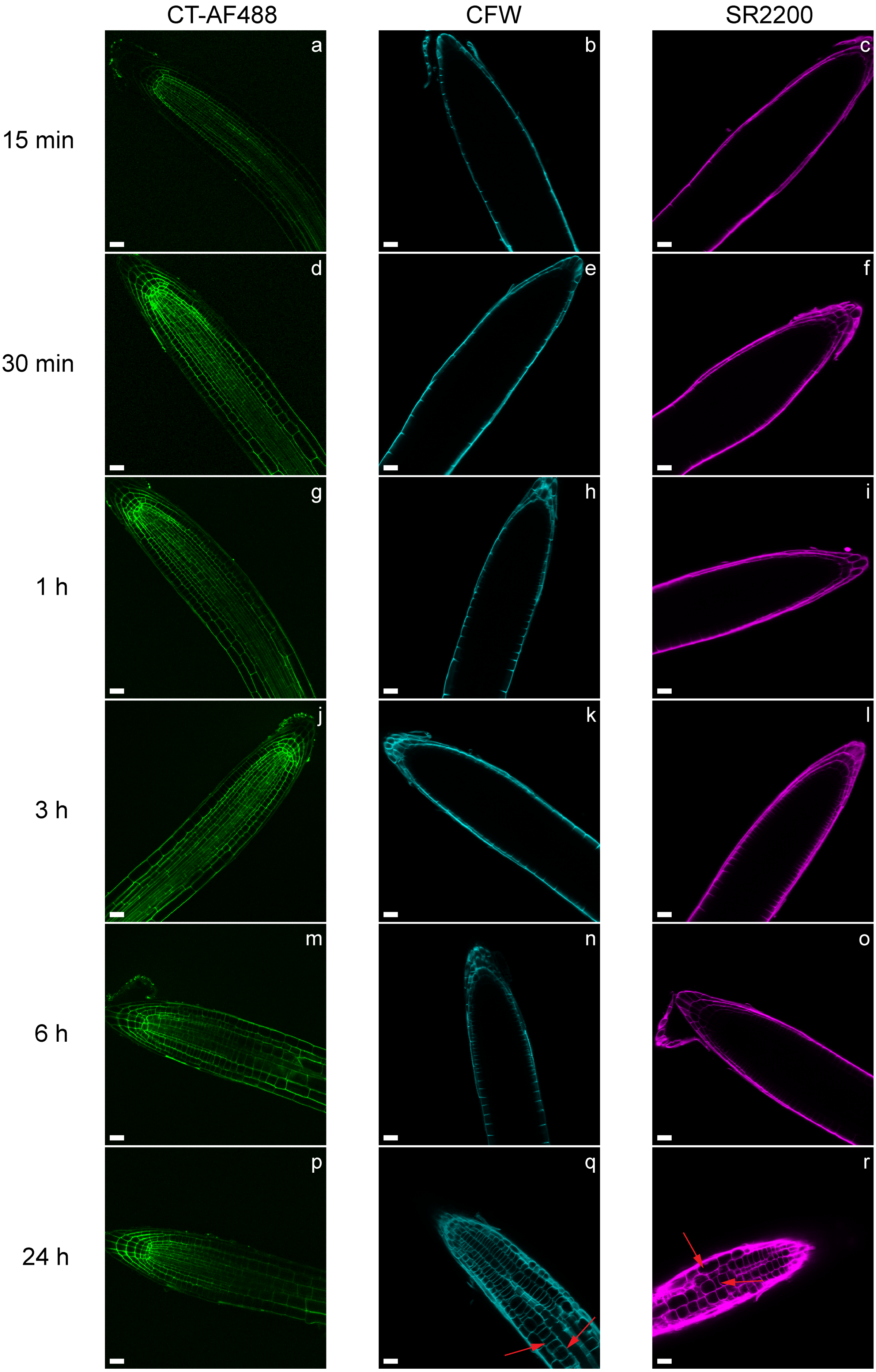


**Figure S7:** penetration of 40 $\mu$M CarboTag-AF488 (CT-AF488, green), 50 ug/ml Calcofluor White (CFW, cyan) and 1x1000 diluted Renaissance 2200 (magenta, SR2200) over time. Red arrows (q,r) indicate altered cell shapes caused by 24 h exposure to CFW and SR2200. Scale bars represent 25 $\mu$m.

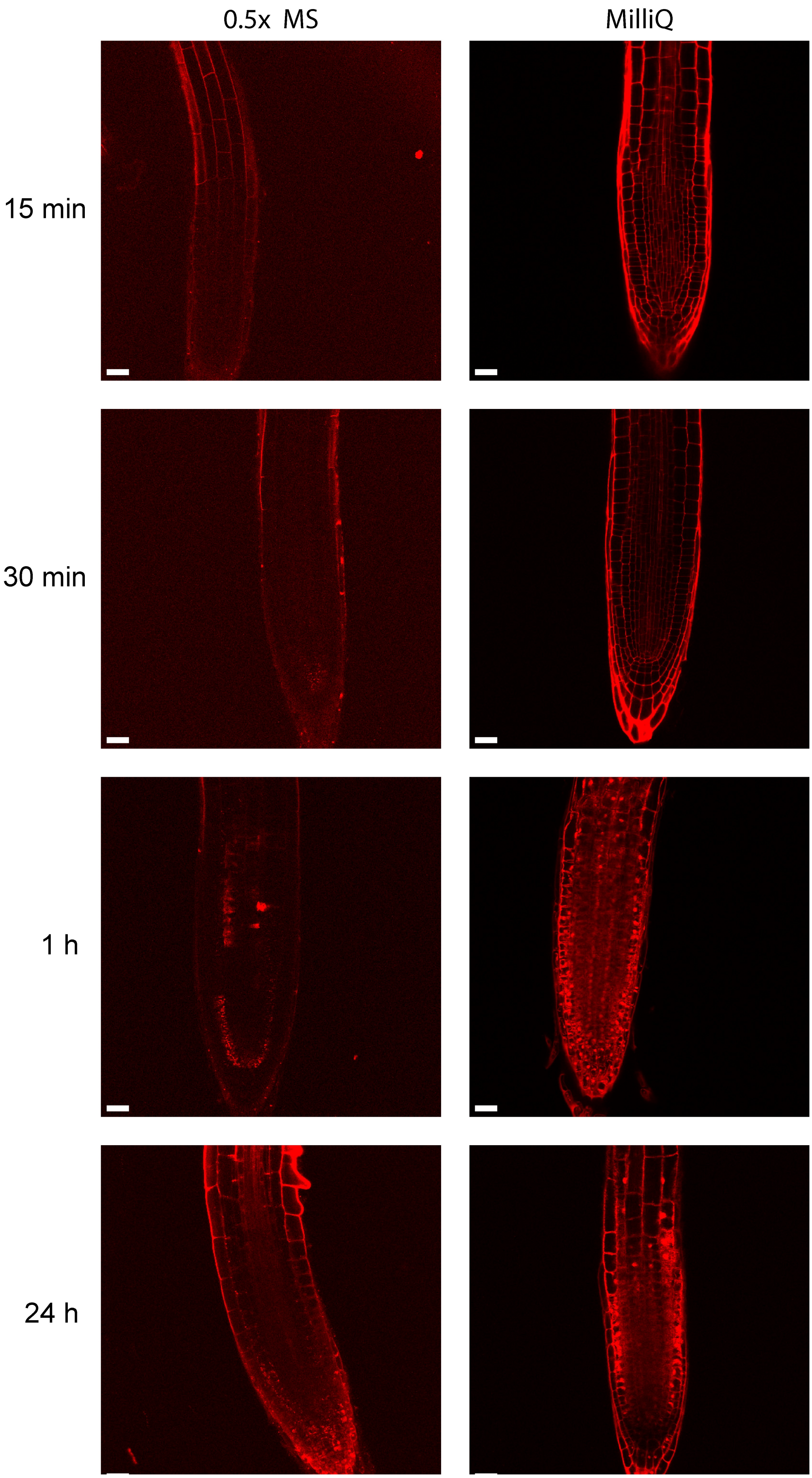


**Figure S8:** penetration of 15 $\mu$M propidium iodide (PI, red) over time in 0.5x MS and MilliQ water. Scale bars represent 25 $\mu$m.

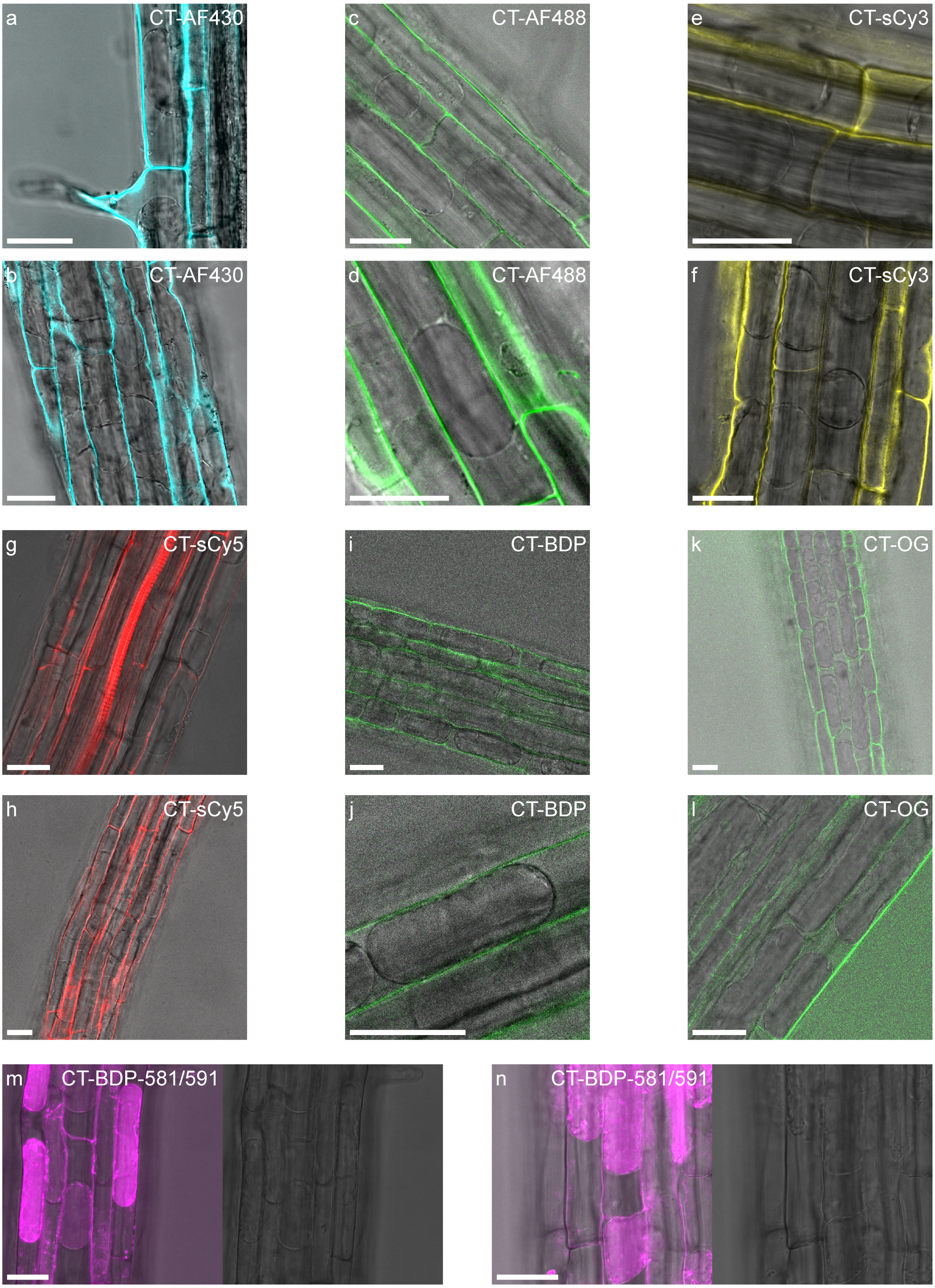


**Figure S9:** plasmolysis of roots incubated with 40$\mu$M CT-AF430 (a,b), CT-AF488 (c,d), CT-sCy3 (e,f), CT-sCy5 (g,h), CT-BDP (i,j), CT-OG (k,l) and 10 $\mu$M CT-Ox (m,n) treated with 0.5 M mannitol. Scale bars represent 25 $\mu$m

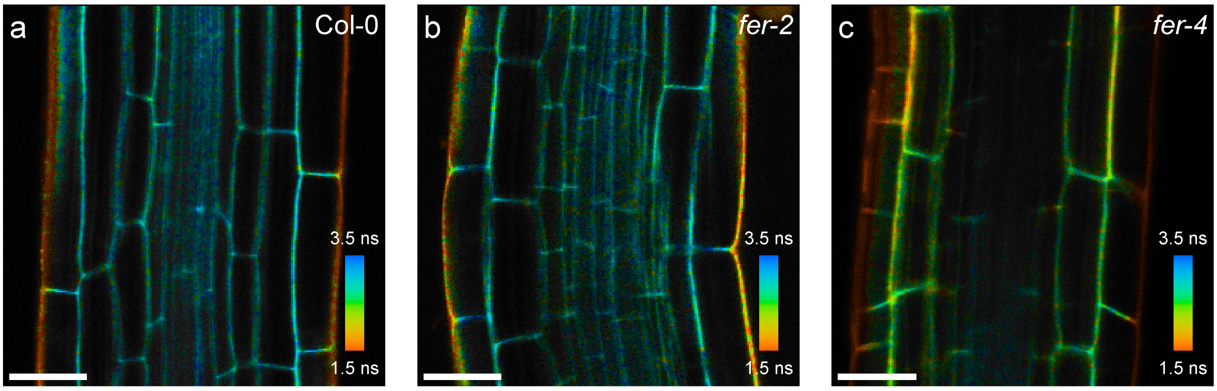


**Figure S10:** FLIM images of Col-0 (a), *fer-2* (b) and *fer-4* (c) mutants incubated with CT-BDP. Scale bars represent 25 $\mu$m.

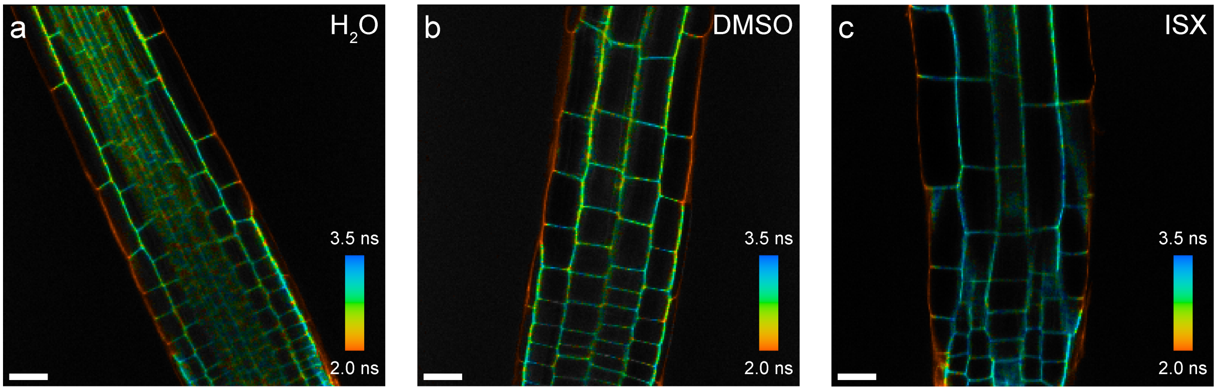


**Figure S11:** FLIM images of non-treated (a), mock treated (b) and isoxaben (ISX) treated (c) Col-0 incubated with CT-BDP. Scale bars represent 25 $\mu$m.

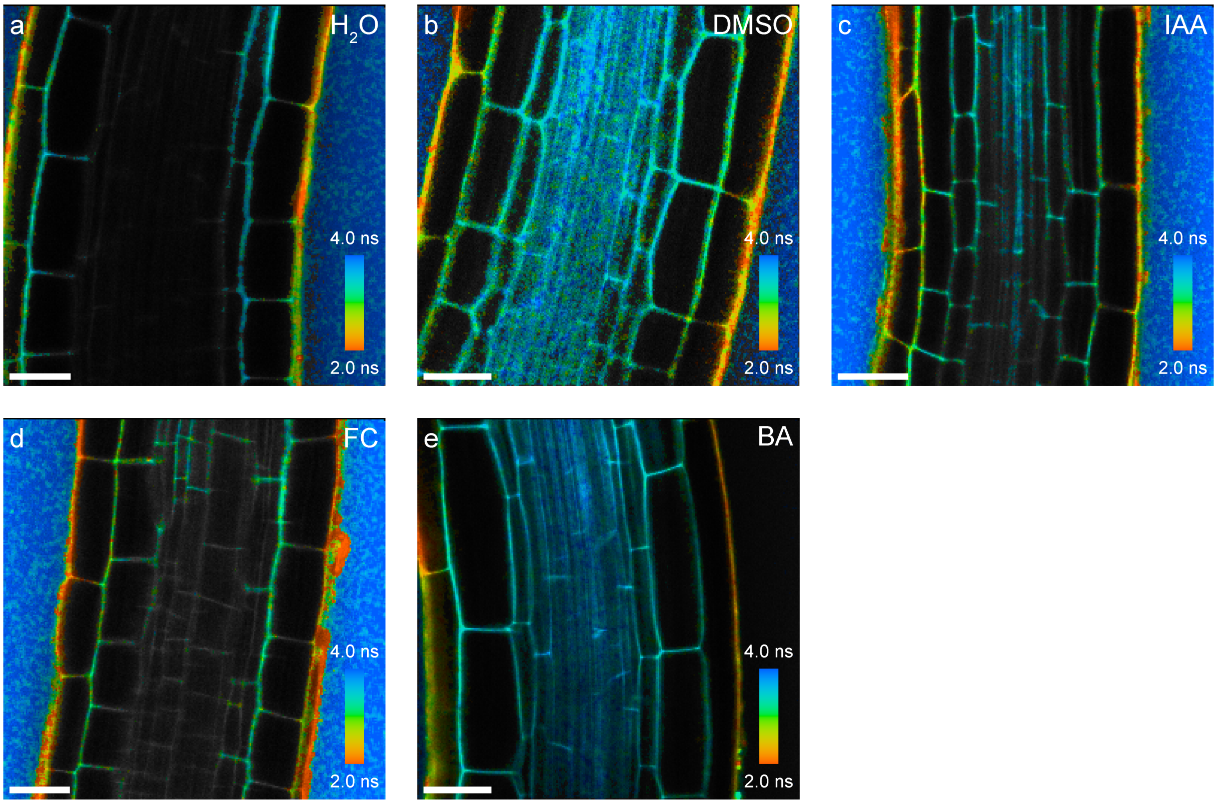


**Figure S12:** FLIM images of roots incubated with 40 $\mu$M CT-OG after 10 minutes of no treatment ($H_2O$, a), mock treatment (DMSO, b), 1 $\mu$M auxin (IAA, c), 10 $\mu$M fusicoccin (FC, d) and 1 $\mu$M benzoic acid (BA, e). Scale bars represent 25 $\mu$m.

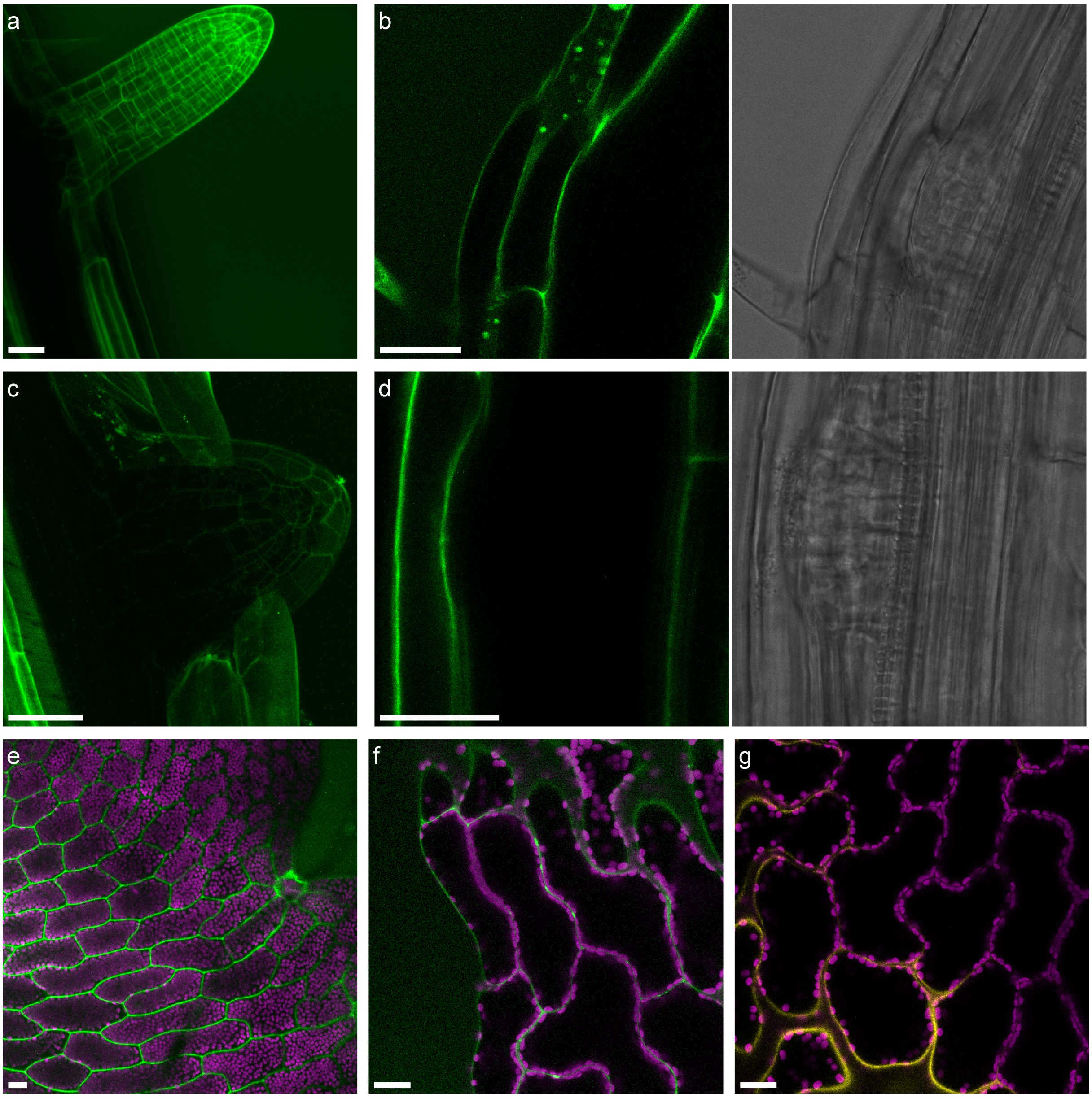


**Figure S13:** a) Average intensity projection of an emerged lateral root stained with CT-AF488 shows good staining. Internal lateral root primordia (b,d) and recently emerged lateral root (c) show no or limited staining. e) Maximum intensity projection of a fern thallus stained with CT-AF488 shows decent staining but is difficult to replicate. f,g) fern thalli stained with CT-AF488 (f) and CT-sCy3 (g) show poor staining even after 6 h. Scale bars represent 25 $\mu$m.

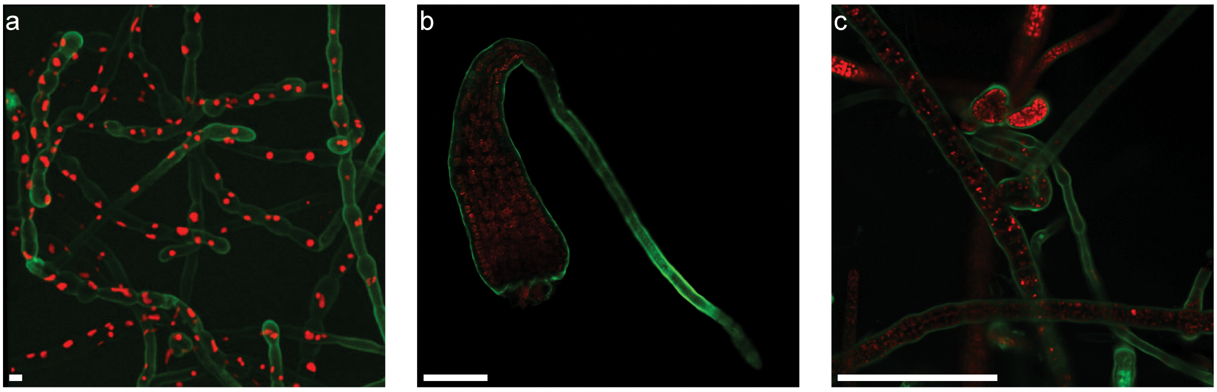


**Figure S14:** Various algae species stained with CT-AF488. *Ectocarpus* (a) and *Fucus serratus* (b) have a clearly highlighted cell wall (green), chloroplast autofluorescence is shown in red. *Sphacelaria* and *Ectocarpus* are clearly distinguishable when stained with CT-AF488. Scale bars represent 1 mm.

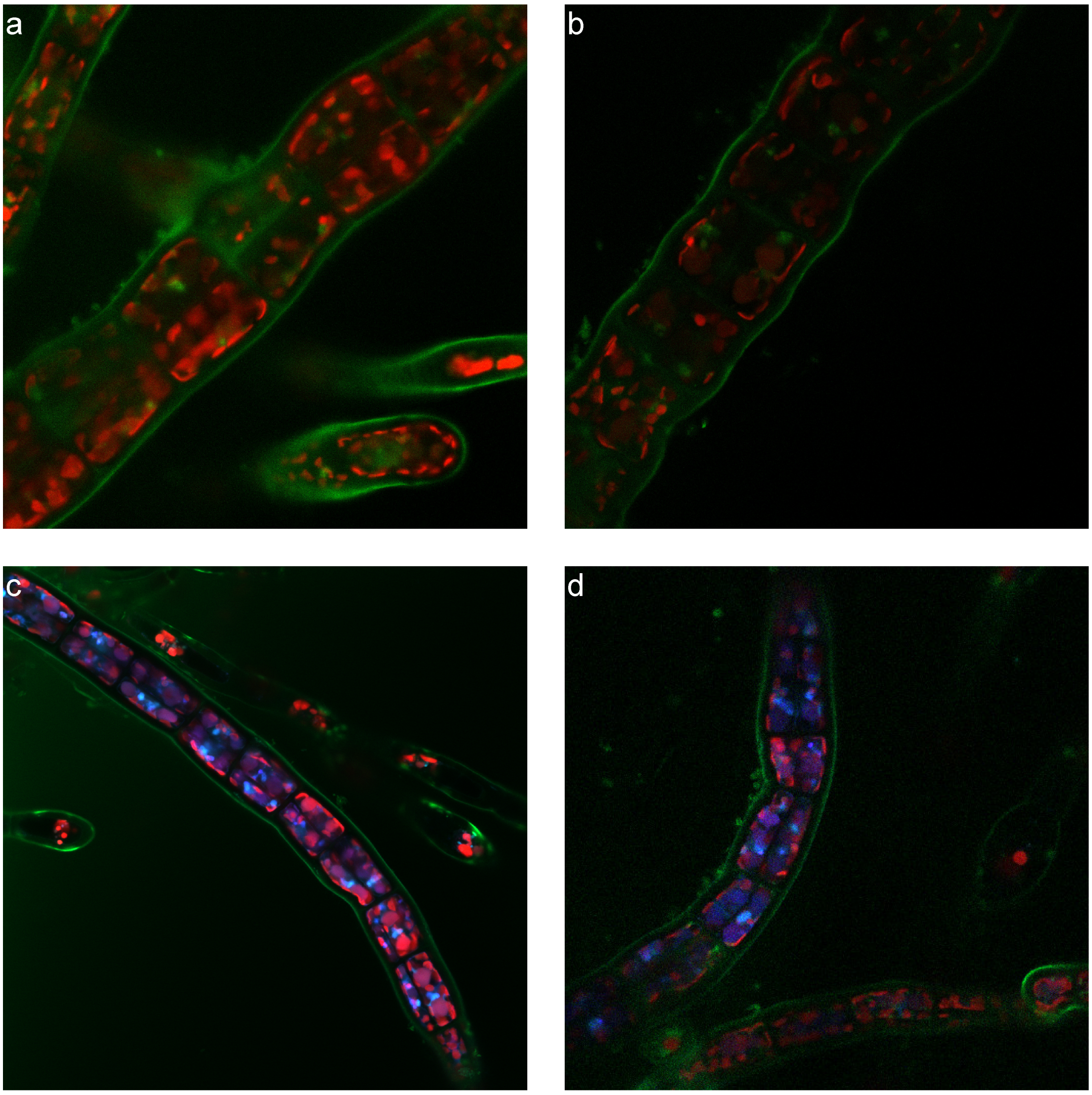


**Figure S15:** a-d) CT-AF488 stained (green) *Sphacelaria* treated with 2.6 M sorbitol. Cell wall shape is maintained after membrane detachment without relocation of CT-AF488 signal, indicating cell wall specificity in this organism. Chloroplast fluorescence is shown in red, non-chloroplast fluorescence in blue.

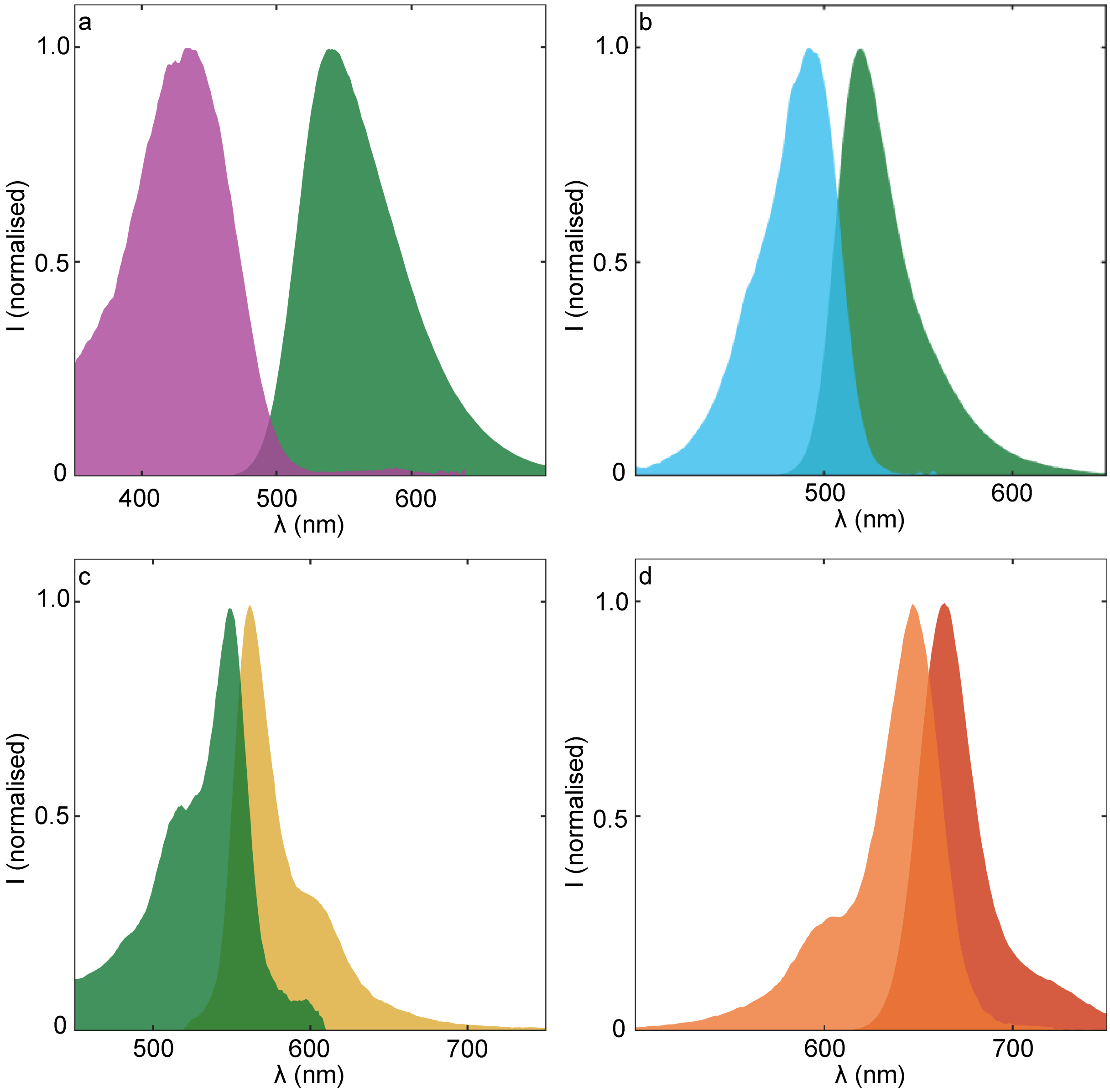


**Figure S16:** excitation emission spectra of CT-AF430 (λ ex = 430 nm, λ em = 650 nm), CT-AF488 λ ex = 450 nm, λ em = 570 nm), CT-sCy3 λ ex = 510 nm, λ em = 620 nm) and CT-sCy5 λ ex = 600 nm, λ em = 710 nm).

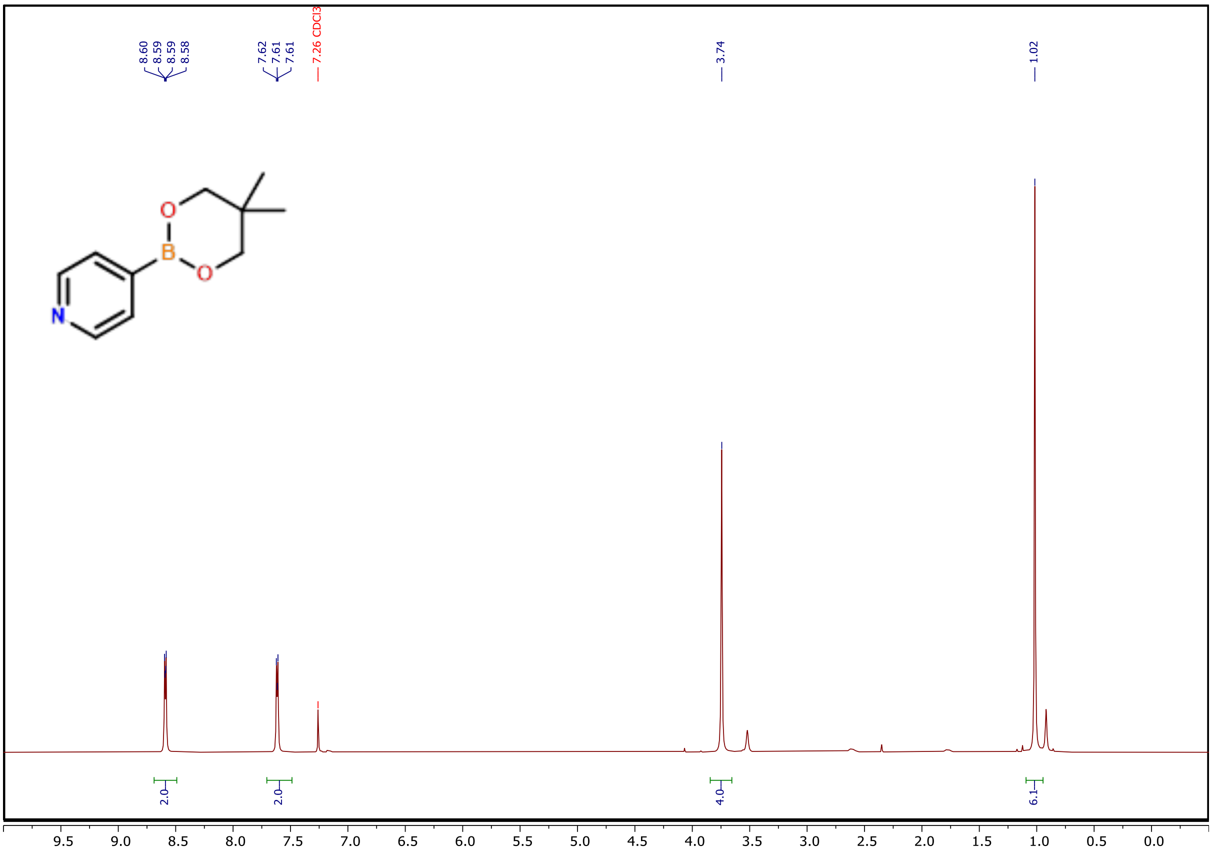


**Figure S17:** $^1$H NMR spectrum of **1**

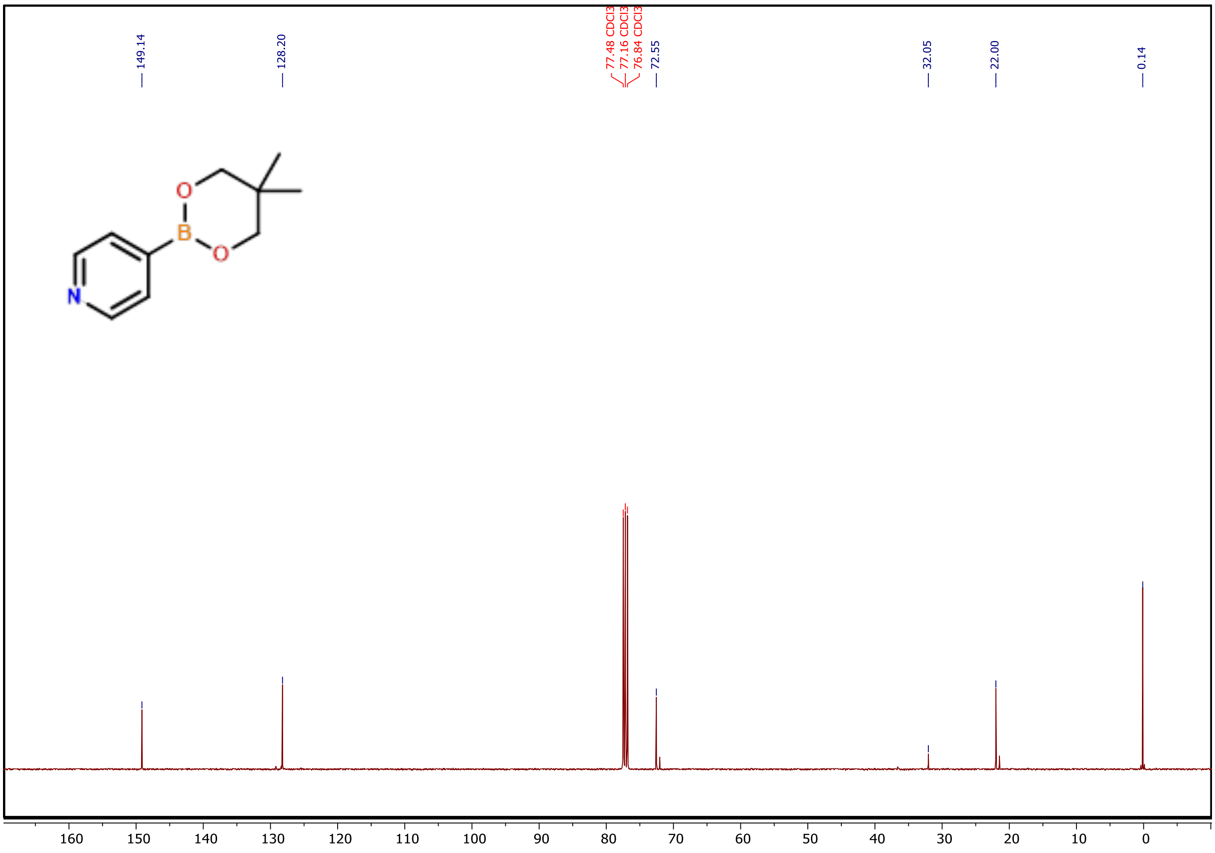


**Figure S18:** $^{13}$C NMR spectrum of **1**

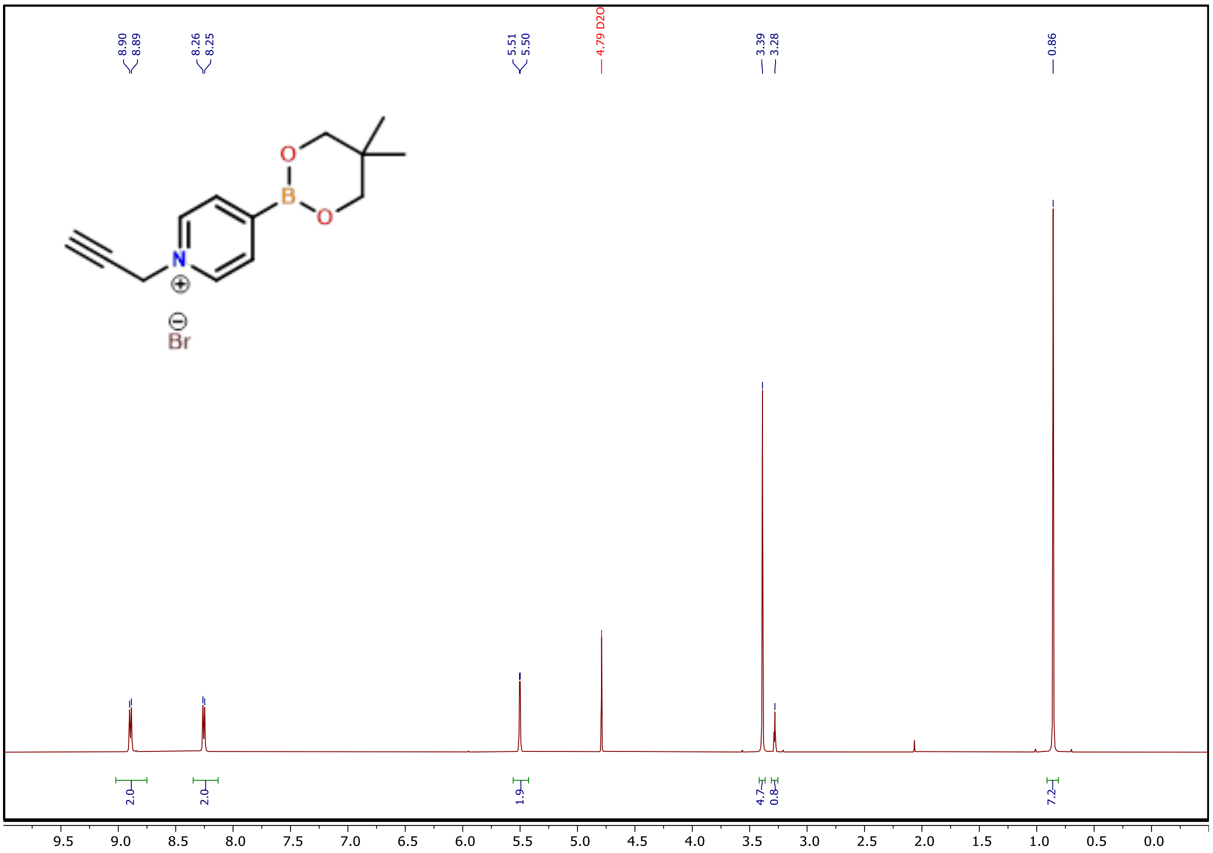


**Figure S19:** $^{1}$H NMR spectrum of **2**

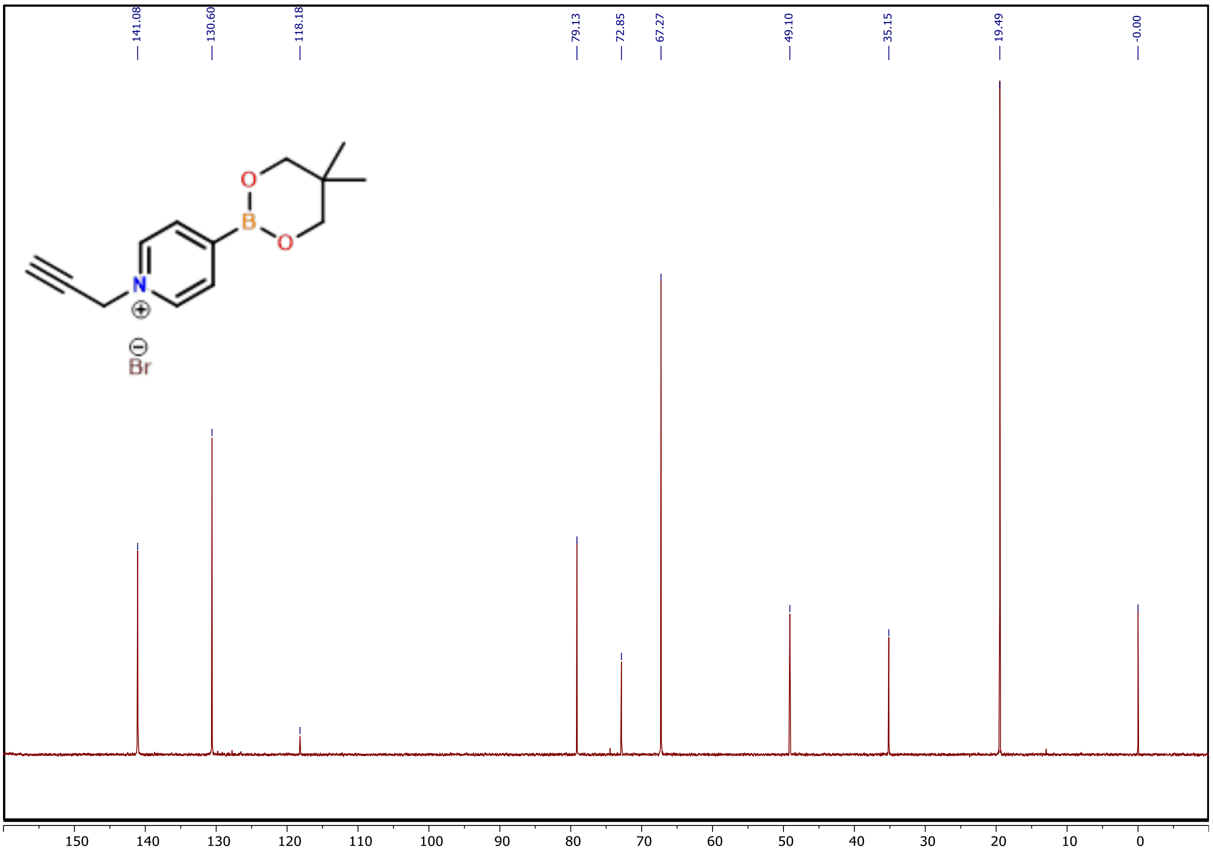


**Figure S20:** $^{13}$C NMR spectrum of **2**

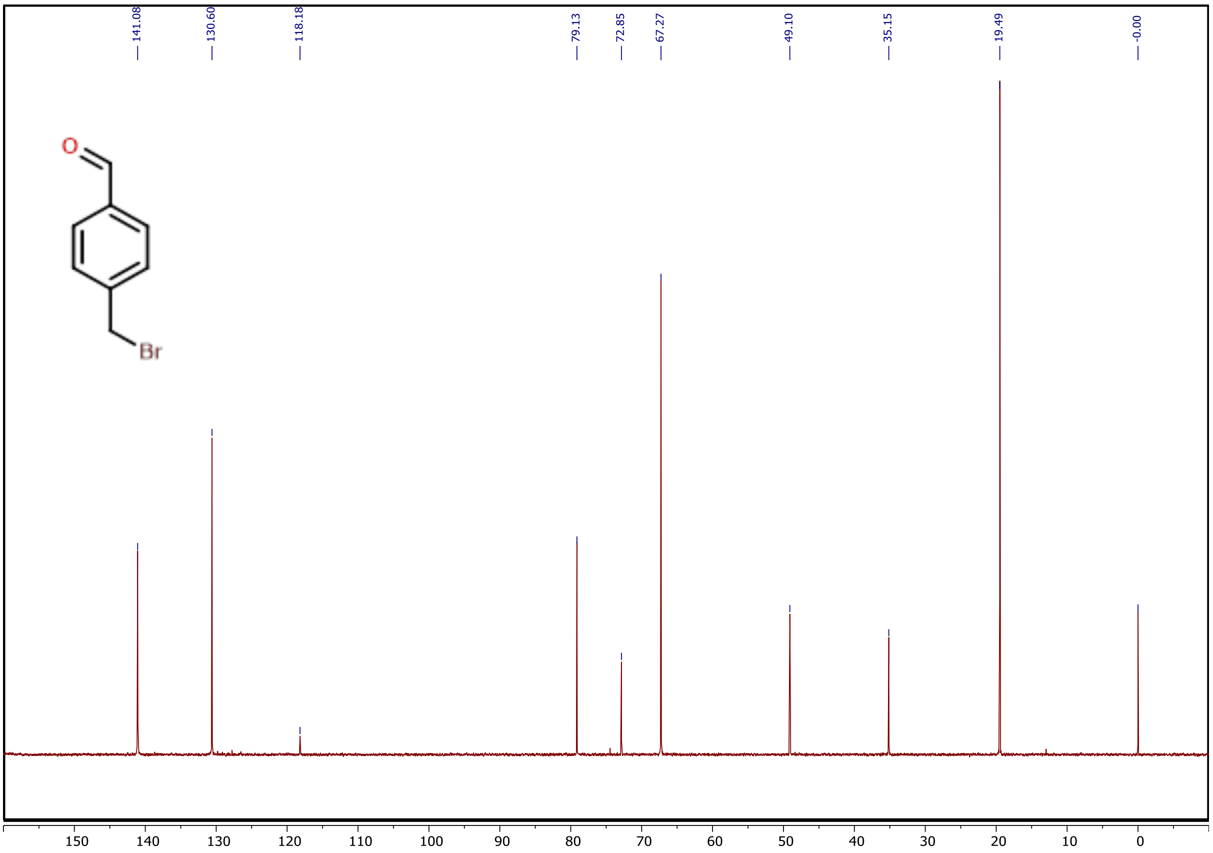


**Figure S21:** $^{1}$H NMR spectrum of **3**

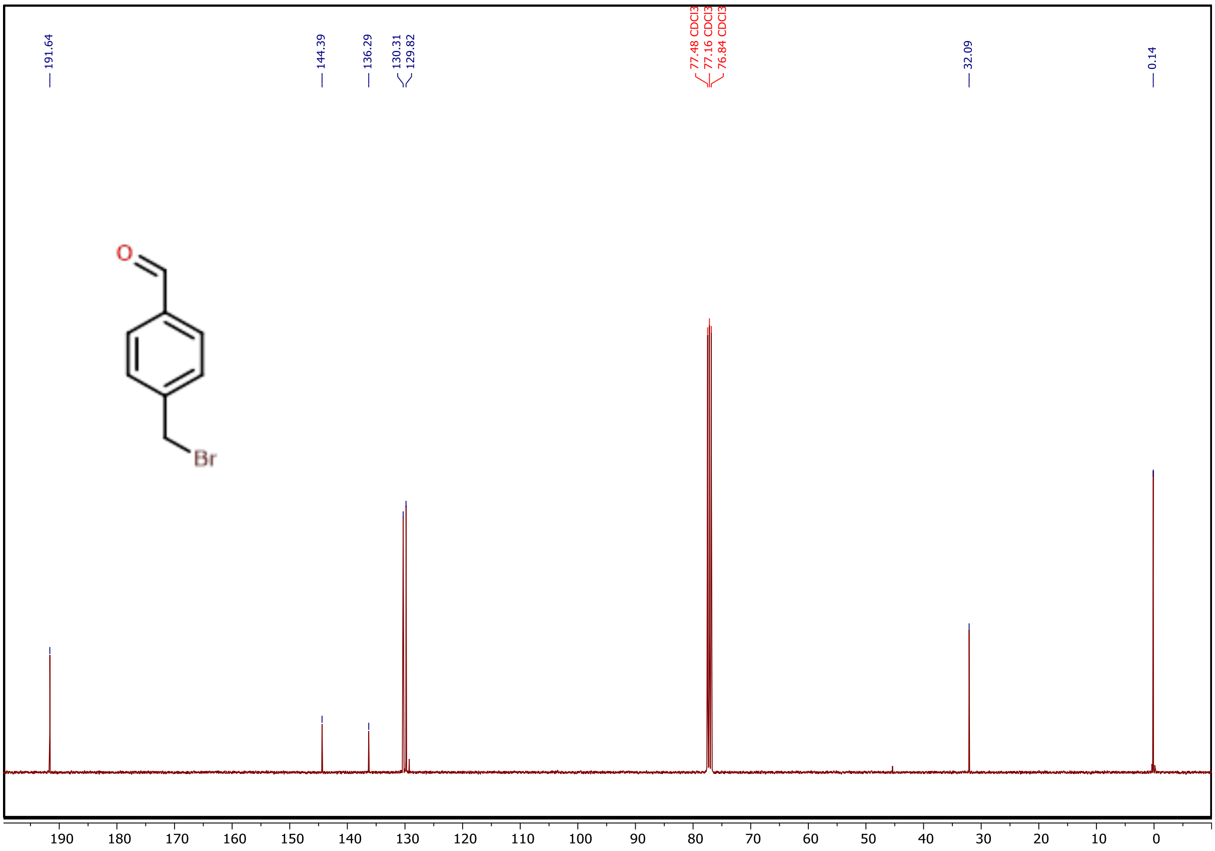


**Figure S22:** $^{13}C$ NMR spectrum of **3**

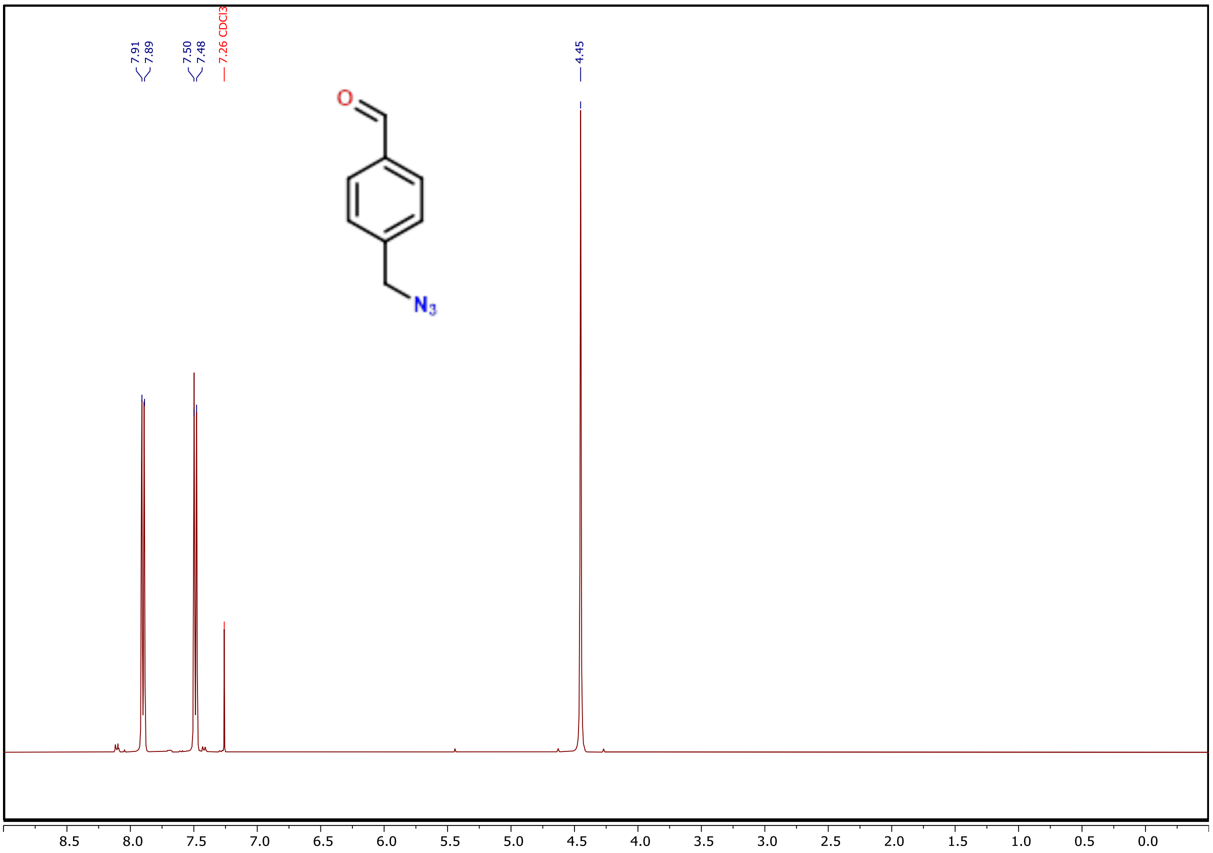


**Figure S23:** $^{1}H$ NMR spectrum of **4**

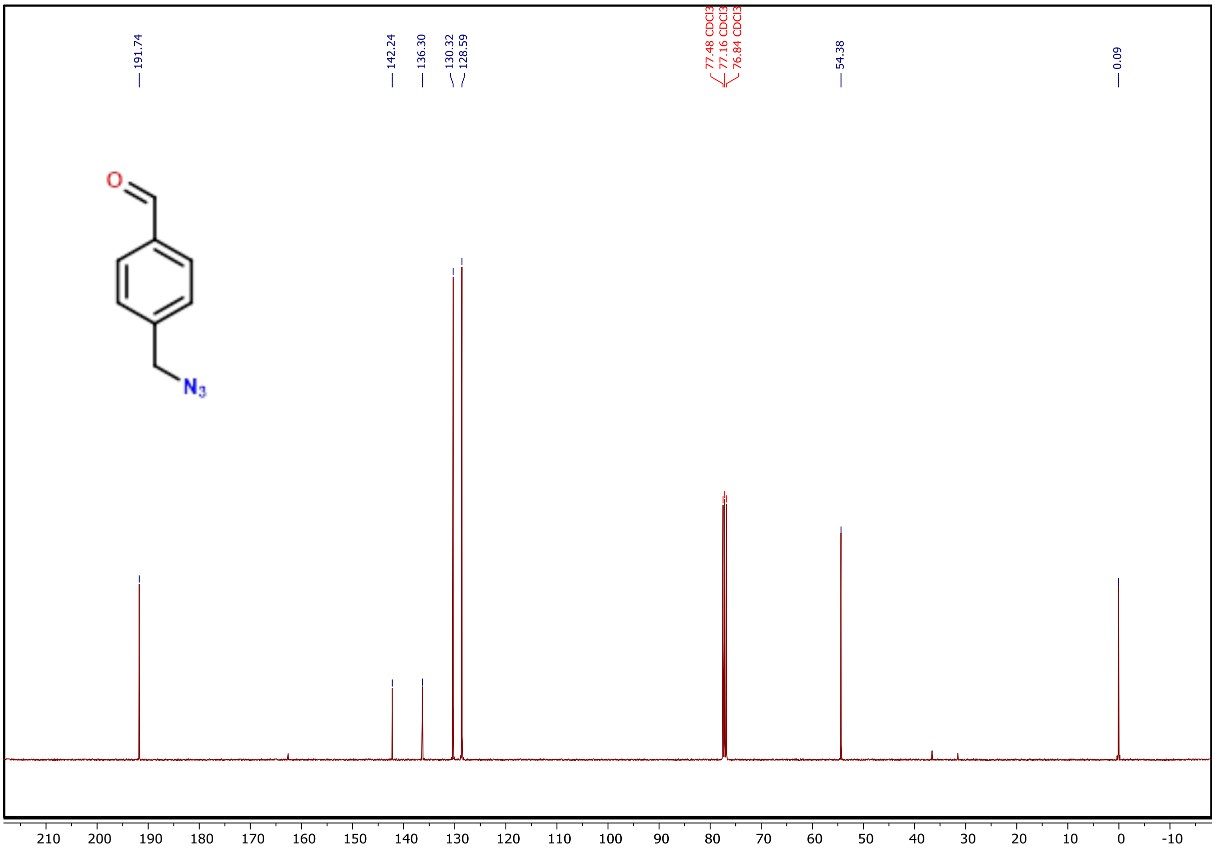


**Figure S24:** $^{13}C$ NMR spectrum of **4**

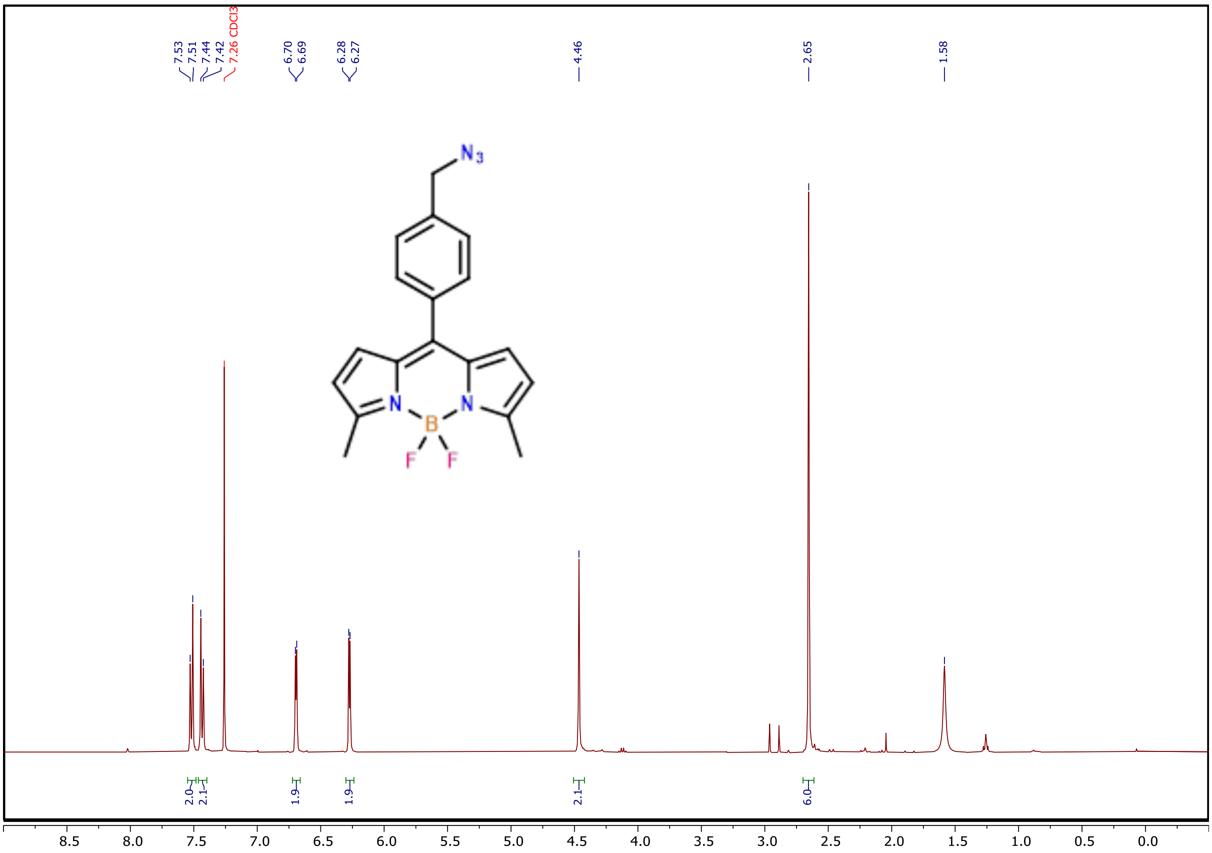


**Figure S25:** $^1$H NMR spectrum of **5**

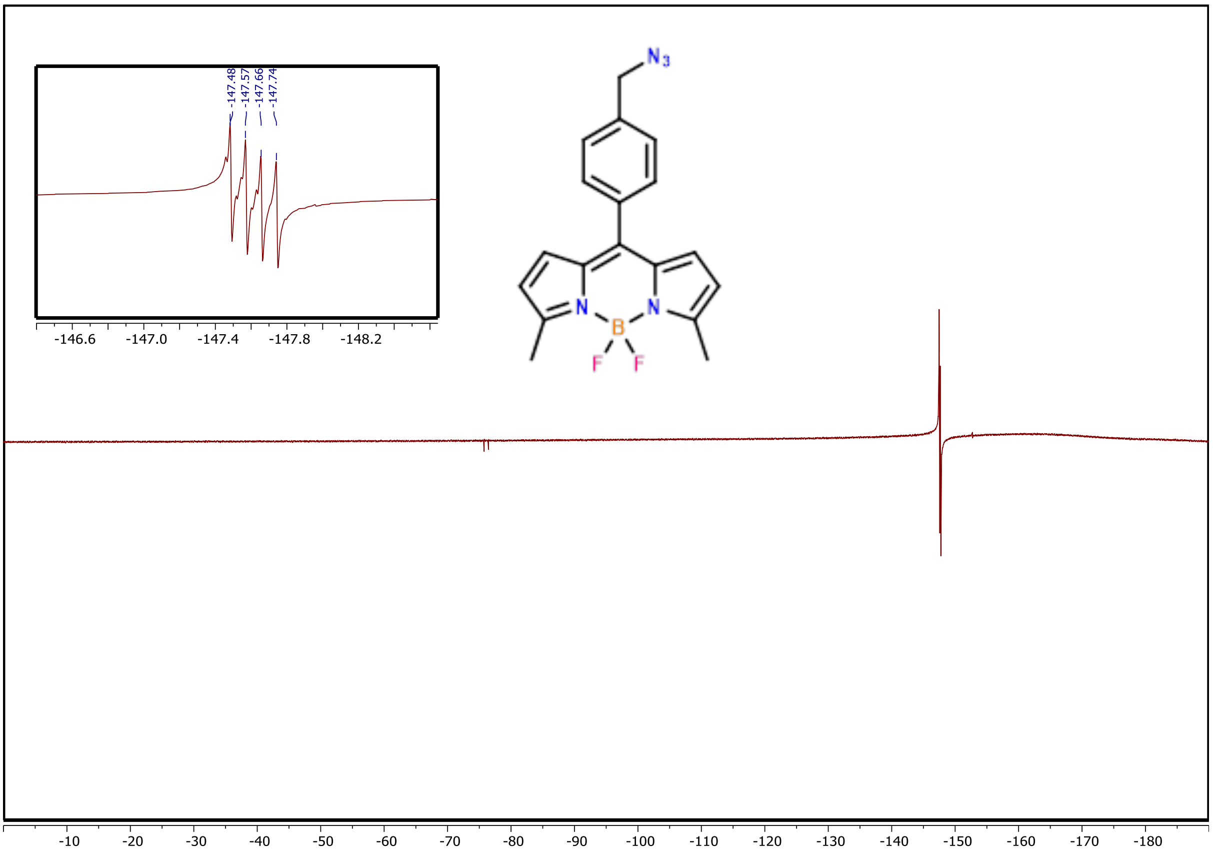


**Figure S26:** $^{19}F$ NMR spectrum of **5**

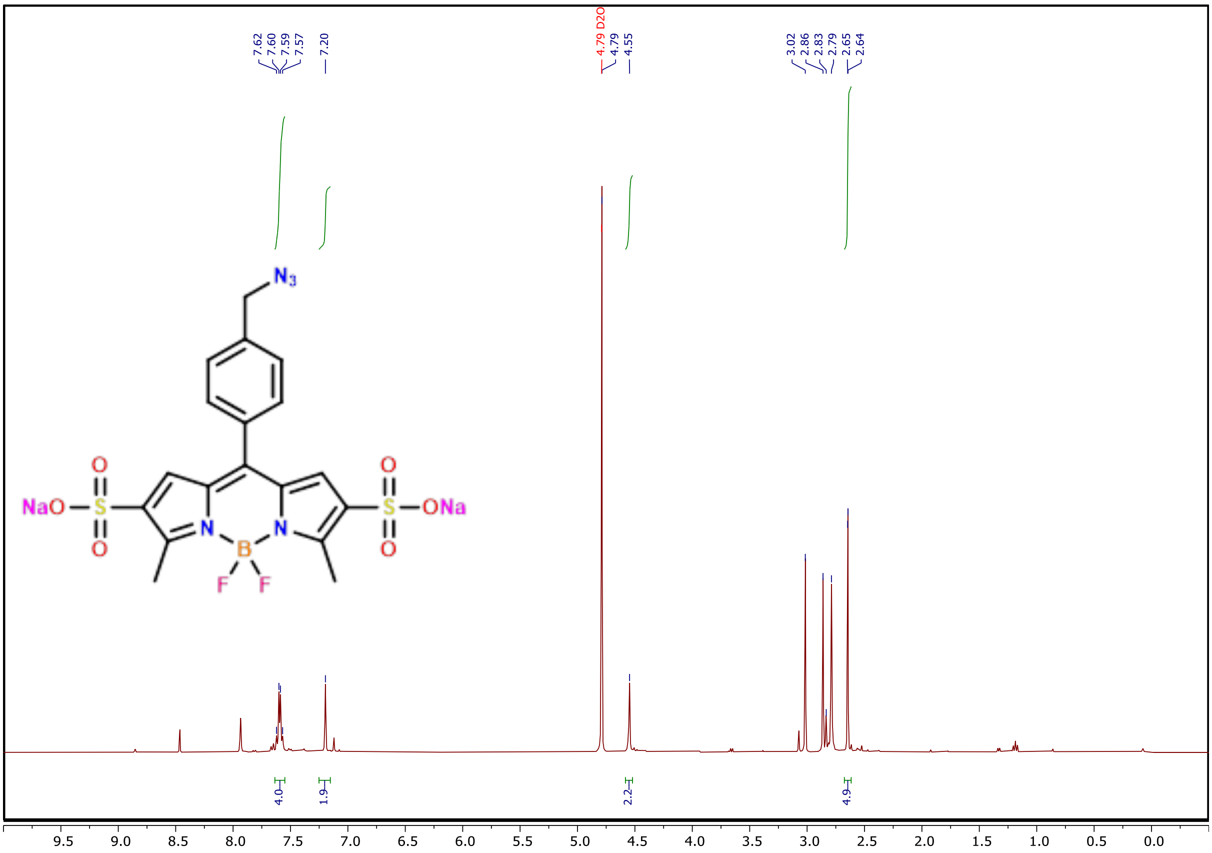


**Figure S27:** $^{1}H$ NMR spectrum of **6**
